# Molecular and functional dissection of the AP2-I/BDP1 transcriptional complex of the malaria parasite reveals a role beyond the red blood cell stages

**DOI:** 10.1101/2025.11.26.690747

**Authors:** Marine Le Berre, Lucille Mellottee, Anagha Rajesh Salvi, Pauline Touarin, Sylvie Auxilien, Virginie Ropars, Marie Chaptal, Cindy Ramiole, Abhinay Ramaprasad, Sylvie Nessler, Joana M. Santos

**Affiliations:** Institute for Integrative Biology of the Cell (I2BC), Paris-Saclay University, CEA, CNRS, 91198 Gif-sur-Yvette, France; Centre for Parasitology, University of Glasgow, School of Infection and Immunity, Sir Graeme Davies Building, 120 University Place, Glasgow, G12 8TA, UK

## Abstract

The transcription factor AP2-I and the bromodomain protein BDP1 of the malaria parasite bind upstream of several genes implicated in red blood cell invasion. Here, we demonstrate that they form a complex and that AP2-I recruits BDP1 to the target genes. The ACDC domain of AP2-I interacts with a BDP1 region called BAAS, located between the ankyrin and the bromodomain. The ankyrin, in turn, binds to a conserved AP2-I specific region called ASAIR. Three of the interacting domains are parasite-specific, allowing inhibition without affecting the host. We further show that AP2-I is essential for asexual blood stage development, playing multiple roles, and that AP2-I and BDP1 function beyond invasion, binding to the promoters of the gametocytogenesis regulators *gdv1* and *ap2-g*, suggesting a role for the AP2-I/BDP1 complex in parasite sexual commitment. Overall, our study highlights the essential and unique nature of the AP2-I/BDP1 complex and its potential as a novel antimalarial drug target.

**Author summary:** The transcription factor AP2-I and the bromodomain protein BDP1 of the malaria parasite have previously been shown to be implicated in red blood cell invasion by the human malaria parasite. Here we show that they form a complex and identify the interacting domains on each protein. Three of these are parasite-specific, allowing inhibition without affecting the human host. We also show that AP2-I is essential for parasite survival in the blood and that the protein is implicated in other processes beyond red blood cell invasion. Namely, AP2-I, but also BDP1, bind to the promoters of the gametocytogenesis regulators *gdv1* and *ap2-g*, suggesting a role for the AP2-I/BDP1 complex in parasite sexual commitment, without which the parasite cannot be transmitted.

## Introduction

Malaria, caused by *Plasmodium* parasites, leads to over half a million deaths each year worldwide (WHO). The most prevalent species are *P. falciparum* and *P. vivax*. The symptomatic phase of malaria is associated with repeated cycles of asexual replication in red blood cells (RBC). Alternatively, a subset of parasites commits to sexual development, differentiating into male and female gametocytes that are transmitted to the mosquito vector.

We previously identified *Pf*AP2-I as the transcription factor activating expression of several genes encoding proteins essential for RBC invasion^1^. AP2-I belongs to the ApiAP2 family, the largest group of DNA-binding proteins in Apicomplexa^2^. It contains three AP2 DNA-binding domains and an N-terminal ACDC domain of unknown function^3^, present only in a subset of ApiAP2 proteins (Fig. 1a).

**Fig 1.**
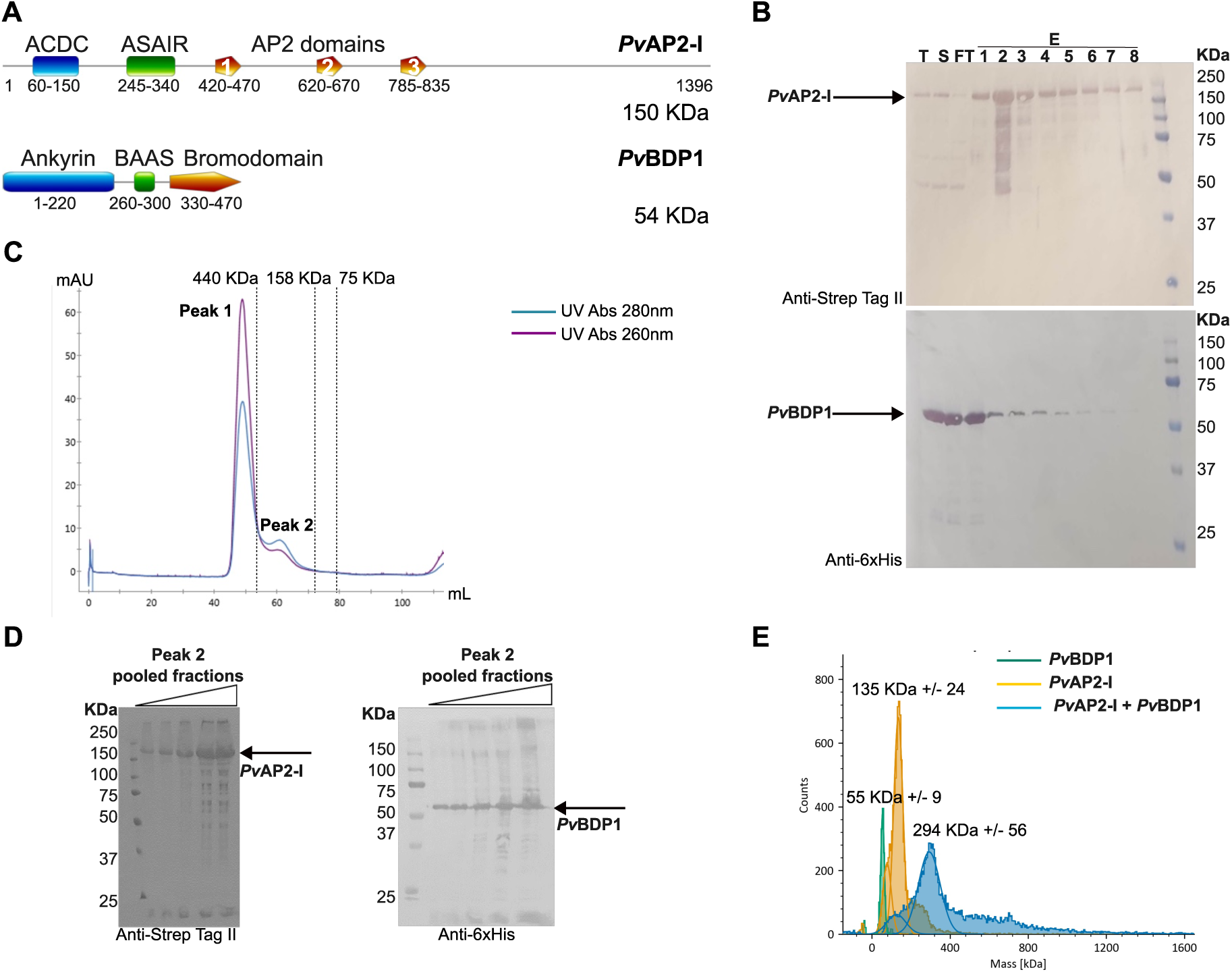
- AP2-I and BPD1 interact directly. **a.** Schematic representation of the *P. vivax* AP2-I and BDP1 proteins, drawn to scale, showing domain organization and protein size. AP2-I contains three AP2 DNA-binding domains (1-3 in orange) as well as an N-terminal ACDC domain (blue), which is only observed in a subset of ApiAP2 proteins. We identify here a previously uncharacterized conserved region that we named ASAIR (AP2-I Specific Ankyrin-Interacting Region; green). BDP1 possesses a C-terminal bromodomain (orange) and an N-terminal ankyrin repeat domain (blue). We named a newly identified conserved region BAAS (Bromodomain-Ankyrin-Associated Sequence; green). **b**. Western blot analysis of elution fractions from the Strep-Trap column following co-expression of Strep-tagged *Pv*AP2-I and His-tagged *Pv*BDP1 in insect cells. Co-elution of both proteins indicates stable complex formation. T: total cell extract, S: soluble fraction, FT: flow-through, E: elution fractions. **c**. Size Exclusion Chromatography (SEC) profile of the purified *Pv*AP2-I/*Pv*BDP1 complex using a Superdex 200 16/60 column. Peak 1 corresponds to high molecular weight DNA-bound aggregates (since the absorbance at 260nm, purple, is higher than at 280nm, blue) eluted in the column’s void volume. Peak 2 corresponds to a ∽300 kDa protein complex. **d**. Western blot analyses of increasing concentrations of the pooled SEC fractions from peak 2 confirmed formation of a stable complex. L: Loaded sample eluted from the Strep-Trap column, E: elution fractions. **e.** Mass photometry analysis of an equimolar mixture of *Pv*AP2-I and *Pv*BDP1 (blue) supports dimerization upon complex assembly, as compared to isolated *Pv*BDP1 (green) and *Pv*AP2-I (yellow).

The bromodomain protein *Pf*BDP1 also regulates transcription of invasion genes^4^. BDP1 comprises an N-terminal ankyrin domain, a central region predicted as intrinsically disordered, and a C-terminal bromodomain (Fig. 1a), recognizing acetylated lysines on histones^4–6^.

Genome-wide mutagenesis indicate that *ap2-i* and *bdp1* are essential^7^. Previous attempts to knockout or knockdown *Pf*AP2-I failed^1,8,9^ but an inducible knockdown (iKD) of *Pf*BDP1 is available^4^. Loss of *Pf*BDP1 impairs RBC invasion^4^, indicating that *Pf*BDP1 is indispensable.

Multiple lines of evidence suggest that *Pf*AP2-I and *Pf*BDP1 form a complex: Both proteins bind the same DNA motif^1,4^; 65% of *Pf*AP2-I target genes, identified by chromatin immunoprecipitation (ChIP), are also bound by *Pf*BDP1^1,4,10^; immunoaffinity purification (IP) identified *Pf*BDP1 as a *Pf*AP2-I interacting protein^1^; and, *Pf*AP2-I binding to a synthetic oligonucleotide containing its DNA-binding motif recruits *Pf*BDP1^11^. *Pf*AP2-I binds to the target genes before *Pf*BDP1^1^, suggesting that *Pf*AP2-I binds DNA first and then recruits *Pf*BDP1 to promote transcription. This model is supported by the observation that although the *Pf*BDP1 bromodomain binds acetylated histones, not all promoters marked by acetylated histones are bound by *Pf*BDP1^4^, indicating that its recruitment to the chromatin depends on additional protein-protein interactions. However, a direct interaction between *Pf*AP2-I and *Pf*BDP1 has never been demonstrated.

*Pf*AP2-I and *Pf*BDP1 may also play roles beyond RBC invasion. Both are expressed in sporozoites^12^, the parasite forms invading liver cells, and in ookinetes^13^, invading the mosquito midgut. *Pf*AP2-I has also been detected in gametocytes^14^ and it interacts with *Pf*AP2-G^15^, the transcription factor triggering sexual commitment. *Pf*AP2-I and *Pf*AP2-G bind together upstream of several invasion genes^15^, but the functional significance of this interaction remains unclear.

Orthologues of *Pf*AP2-I and *Pf*BDP1 are encoded in *P. vivax*^16^, suggesting a conserved role for the proteins. AP2-I represents a promising drug target because it is essential, it lacks human homologues, it is conserved across the genus, and it is expressed at multiple stages of the parasite life cycle. Moreover, recent studies demonstrated the druggability of ApiAP2 proteins^17^ and structural modelling suggests that the bromodomain of BDP1 is sufficiently distinct from other proteins to enable selective inhibition^5^.

In this study, we performed an in-depth investigation of AP2-I and BDP1, as well as their interaction, both *in vitro* and in *P. falciparum*. We demonstrate that the two proteins directly interact and identify interacting domains. In parasites, we confirm that *Pf*BDP1 associates with *Pf*AP2-I. We also show that *Pf*AP2-I is essential for parasite growth. Finally, we provide evidence that the two proteins participate in sexual commitment. Together, our results highlight the essential and unique nature of the AP2-I/BDP1 complex in *Plasmodium* and underscore its potential as a novel antimalarial drug target.

## Results

### AP2-I and BDP1 interact directly

The *in vitro* characterization of AP2-I and BDP1 was performed using the *P. vivax* orthologues, because *Pv*AP2-I contains shorter unstructured poly-asparagines stretches than *Pf*AP2-I, which facilitates heterologous expression. Strep-tagged *Pv*AP2-I (151 kDa) and His-tagged *Pv*BDP1 (56 kDa) were co-expressed in insect cells, and the soluble protein fraction was applied to a Strep-Trap affinity column. Western-blot analysis of the eluted fraction demonstrated that His-tagged *Pv*BDP1 co-eluted with Strep-tagged *Pv*AP2-I (Fig. 1b). His-tagged *Pv*BDP1 was not retained on the Strep-Trap column in the absence of *Pv*AP2-I (data not shown), indicating that *Pv*AP2-I and *Pv*BDP1 directly interact. Complex formation was also observed when the two proteins were expressed separately. His-tagged *Pv*BDP1 expressed in *E. coli* co-eluted with Strep-tagged *Pv*AP2-I purified from insect cells (Supplementary Fig. 1a).

The purified *Pv*AP2-I/*Pv*BDP1 complex was further characterized using size exclusion chromatography (SEC). A major peak containing DNA eluted in the void volume of the Superdex 200 column (Fig. 1c). Western blot analysis revealed that the second peak contained both *Pv*AP2-I and *Pv*BDP1 (Fig. 1d). This peak eluted at a volume corresponding to an estimated molecular mass of ∼330 kDa (Fig. 1c), exceeding the expected weight of a 1:1 heterodimer (207 kDa). Consistent with this result, mass photometry analysis of an equimolar mix of the two proteins revealed a predominant species of 294 ± 56 kDa whereas the individual *Pv*AP2-I and *Pv*BDP1 proteins were monomeric in solution (Fig. 1e). These results suggest that complex formation induces AP2-I dimerization but it is still unclear whether it binds to one or two BDP1 molecules.

### Identification of the AP2-I region recognized by the ankyrin domain of BDP1

To identify the interacting domains of AP2-I and BDP1, various protein fragments were cloned and expressed in *E. coli*, tagged with a C-terminal StrepTag II (*Pv*AP2-I) or a C-terminal His-tag (*Pv*BDP1). Co-purification assays were performed on a Strep-Trap affinity column, and elution fractions were analyzed by Western blot. Complex formation was deduced from co-elution.

The *Pv*BDP1 ankyrin domain (residues 1-220) interacted with full-length *Pv*AP2-I (Fig. 2a), whereas the bromodomain (residues 330-470 did not (data not shown). Interaction was also observed between the *Pv*BDP1 ankyrin domain and the *Pv*AP2-I N-terminal region (residues 1-340) (Fig. 2b, Supplementary Fig. 2a), containing the ACDC domain (residues 60-150) and a conserved sequence (residues 250-340) specific to AP2-I orthologs (Supplementary Fig. 2b).

**Fig 2.**
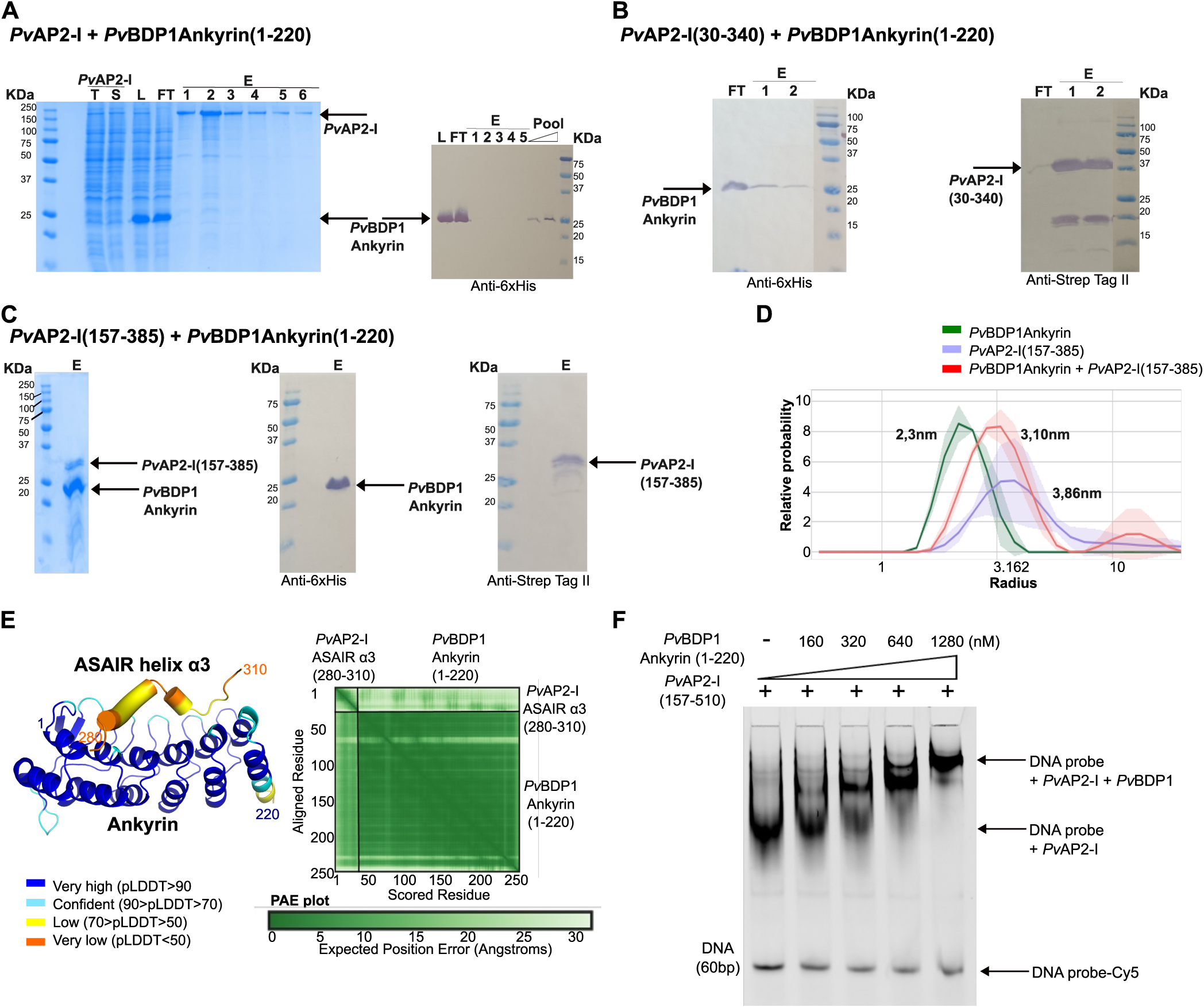
- Identification of the AP2-I region recognized by the ankyrin domain of BDP1. **a**. Co-purification on a Step-Trap column of a mixture of full-length Strep-tagged *Pv*AP2-I expressed in insect cells and His-tagged *Pv*BDP1 ankyrin domain (fragment 1-220, 26 kDa) expressed in *E. coli*. SDS-PAGE and Western-blot blot analyses of the Strep-Trap elution fractions confirm co-elution of the ankyrin domain with *Pv*AP2-I, indicating direct interaction. T: total insect cells extract, S: soluble fraction from insect cells extract, L: loaded mixture, FT: flow-through, E: elution fractions, Pool: increasing concentrations of the elution pool. **b**. Co-purification of the His-tagged *Pv*BDP1 ankyrin domain with the Strep-tagged N-terminal fragment of *Pv*AP2-I (residues 1-340, 39 kDa) co-expressed in *E. coli*. Western blot analysis of the elution fraction demonstrates co-purification of both proteins, suggesting that the N-terminal region of *Pv*AP2-I interacts with the ankyrin domain. A ∼20 kDa protein detected with the anti-Strep Tag antibody corresponds to degradation of the *Pv*AP2-I fragment. FT: flow-through, 1-2: elution fractions. A non-cropped image is shown on Supplementary Fig. 2a. **c.** Co-elution of a Strep-tagged fragment of *Pv*AP2-I (residues 157-385, 26 kDa), containing the AP2-I specific conserved region (residues 250-340), with the His-tagged ankyrin domain of *Pv*BDP1, expressed separately in *E. coli*. SDS-PAGE and Western-blot analyses of the elution fraction (E) confirm a specific interaction, indicating that residues 157-385 of *Pv*AP2-I are sufficient to bind the ankyrin domain. **d.** Dynamic Light Scattering (DLS) analysis of the complex between the ankyrin domain of *Pv*BDP1 and the *Pv*AP2-I (157-385) fragment. DLS results reveal an intermediate hydrodynamic radius for the complex (red) compared to the isolated ankyrin domain (green) or the *Pv*AP2-I fragment (purple), suggesting that binding induces a transition of the *Pv*AP2-I fragment from an unfolded to a partially folded state. **e**. AlphaFold3 model (Rank 1) of the complex between the ankyrin domain of *Pv*BDP1, shown as cartoon, and the ASAIR region of *Pv*AP2-I, encompassing the best predicted α3 helix (residues 280-310) shown as a cylindrical helix. Both are colored according to the pLDDT confidence score of the models. The reliability of the proposed interaction mode is illustrated by the Predicted Aligned Error (PAE) plot. **f.** Electro Mobility Shift Assay (EMSA) showing that high concentrations of the *Pv*BDP1 ankyrin domain alone do not bind DNA but induce a supershift in the mobility of a 60 bp oligonucleotide containing the DNA-binding motif of the first AP2 domain of AP2-I, bound to a fixed amount (320 nM) of the *Pv*AP2-I fragment (157-510), encompassing ASAIR and the first AP2 domain. A non-cropped gel is shown in Supplementary Figure 2h.

To delineate the *Pv*AP2-I region interacting with the *Pv*BDP1 ankyrin domain, we expressed *Pv*AP2-I fragments containing either the ACDC domain or the conserved sequence (residues 30-180 and 157-385, respectively). Whereas full-length *Pv*BDP1 co-eluted with both fragments (Supplementary Figs. 2c-d), the isolated ankyrin domain interacted only with the 157-385 fragment (Fig. 2c and data not shown). Dynamic Light Scattering (DLS) measurements revealed a hydrodynamic radius (R_h_) of 2.3 ± 0.17 nm for the ankyrin domain, 3.86 ± 0.36 nm for the *Pv*AP2-I (157-385) fragment, and an intermediate value of 3.1 ± 0.16 nm for the two proteins mixture (Fig. 2d). These results suggest that the *Pv*AP2-I fragment adopts a more compact conformation upon binding to the ankyrin domain, potentially reflecting a transition from an unfolded to a partially folded state.

The *Pv*AP2-I conserved sequence recognized by the ankyrin domain was named ASAIR for AP2-I Specific Ankyrin-Interacting Region. Secondary structure predictions (Supplementary Fig. 2e) and AlphaFold3 modeling suggested that ASAIR comprises five α-helices, although AlphaFold3 failed to predict a consistent 3D fold (Supplementary Fig. 2f). Nevertheless, a robust model for the complex between the concave face of the *Pv*BDP1 ankyrin domain and the α3 helix of ASAIR (residues 280-310) was obtained (Fig. 2e). This model is highly consistent across all five AlphaFold3 predictions, with a root-mean-square deviation (RMSD) of 0.76 Å and predicted Template Modeling (pTM) and interface pTM (ipTM) scores of 0.84 and 0.34, respectively.

To experimentally validate this interaction, thermal shift assays were performed with a synthetic peptide corresponding to the α3 helix of ASAIR (residues 280-310). The melting temperature (Tm) of the *Pv*BDP1 ankyrin domain increased from 50.0°C to 50.7°C or 51.2°C upon addition of a 1:10 or a 1:40 molar ratio of the peptide, respectively (Supplementary Fig. 2g). No thermal shift was observed with the isolated peptide (data not shown), indicating that stabilization of the *Pv*BDP1 ankyrin domain resulted from specific interaction with the ASAIR peptide.

Finally, to test whether the *Pv*BDP1 ankyrin domain can interact with ASAIR when AP2-I is bound to DNA, Electrophoretic Mobility Shift Assays (EMSA) were performed with a *Pv*AP2-I fragment encompassing both ASAIR and the first AP2 DNA-binding domain (residues 157-510) using a 60-bp probe contained the GTCGAC motif specifically recognized by the first AP2 domain of AP2-I^18^. Whereas the ankyrin domain alone did not bind the DNA probe (Supplementary Fig. 2h), addition of increasing concentrations of the ankyrin domain (fragment 1-220) resulted in a super-shift of the DNA-protein complex (Fig. 2f). The ankyrin-ASAIR interaction is thus maintained when AP2-I is bound to DNA.

### Identification of the BDP1 region recognized by the ACDC domain of AP2-I

As mentioned above, the *Pv*AP2-I ACDC domain interacted with full-length *Pv*BDP1 (Supplementary Fig. 2c) but not with the isolated ankyrin domain (fragment 1-220) (data not shown), indicating that ACDC recognizes a different region of *Pv*BDP1. Since the *Pv*BDP1 bromodomain (residues 310-470) also did not interact with full-length *Pv*AP2-I (data not shown), the *Pv*BDP1 central region (residues 219-312), predicted as disordered, likely mediates this interaction. Although the central region could not be purified alone, fusions with either the ankyrin domain (fragment 1-312) or the bromodomain (fragment 219-470), co-eluted with the *Pv*AP2-I ACDC domain (Figs. 3a-b), indicating that ACDC interacts with this *Pv*BDP1 region.

**Fig 3.**
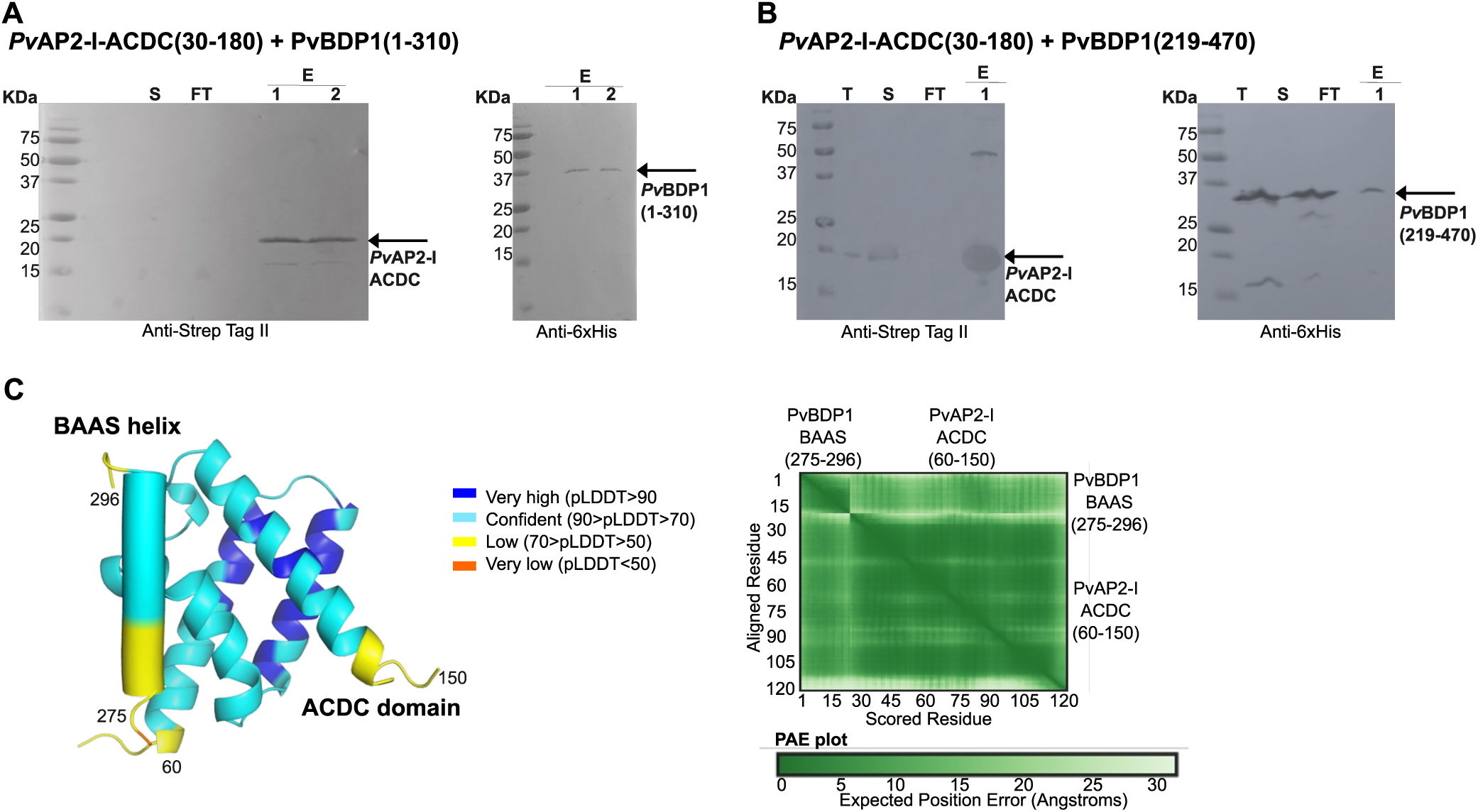
- Identification of the BDP1 region recognized by the ACDC domain of AP2-I. **a**-**b**. Co-purification on a Strep-Trap column of the Strep-tagged ACDC domain of *Pv*AP2-I (residues 30-180, 18 kDa) with the His-tagged central region of *Pv*BDP1 fused to (**a**) the ankyrin domain (fragment 1-310, 36kDa) or (**b**) to the *Pv*BDP1 bromodomain (fragment 219-470, 28 kDa). Western blot analyses of both elution fractions demonstrate that the ACDC domain interacts with the central region of *Pv*BDP1. S: soluble fraction, FT: flow-through, E: elution fractions. **c**. AlphaFold3 predicted model (Rank 1) of the complex between the ACDC domain of *Pv*AP2-I (residues 60-150) and the BAAS region of *Pv*BDP1 containing the predicted α helix (residues 275-296). The ACDC domain is shown as cartoon and the BAAS helix as a cylinder. The structures are colored according to the pLDDT confidence score. The predicted Aligned Error (PAE) plot supports the reliability of the predicted interaction mode.

In the central region, residues 260-300 are highly conserved among BDP1 orthologues (Supplementary Fig. 3a). We named this conserved segment BAAS (Bromodomain-Ankyrin-Associated Sequence), given its position between the ankyrin domain and the bromodomain. BAAS is only present in Apicomplexa BDP1 proteins, despite the fact that some Alveolata encode proteins containing an ankyrin and a bromodomain (Supplementary Figs. 3b-c).

According to AlphaFold3, BAAS is disordered, except for residues 283-292, which are predicted to form an α-helix (Supplementary Figs. 3d-e). AlphaFold3 also generated a high-confidence model of the BAAS α-helix bound to the *Pv*AP2-I ACDC domain (pTM = 0,72, ipTM = 0.52) (Fig. 3c), with all five predictions converging to an RMSD of 0.69 Å, strongly supporting the predicted interface.

Thermal shift assays further validated this interaction. Addition of a synthetic peptide corresponding to *Pv*BDP1 BAAS (residues 275-296) to the ACDC domain reduced its melting temperature from 61.1°C to 60.1°C or 59.8°C at a 1:10 or 1:40 molar ratio, respectively (Supplementary Fig. 3f). No thermal shift was observed for the peptide alone (data not shown), confirming a direct and specific interaction between *Pv*AP2-I ACDC and *Pv*BDP1 BAAS.

### The AP2-I/BDP1 complex assembles in the parasite after DNA-binding

Immunoprecipitation (IP) of *Pf*AP2-I previously identified *Pf*BDP1 as a partner protein^1^, however reciprocal IP experiments performed by us (Supplementary Table 1) or others^4,10^ with *Pf*BDP1 failed to recover *Pf*AP2-I. These results are consistent with our hypothesis that *Pf*AP2-I recruits *Pf*BDP1 to the chromatin and that only chromatin-bound *Pf*BDP1 associates with *Pf*AP2-I. To identify the protein partners of chromatin-bound *Pf*BDP1, we performed chromatin immunoprecipitation followed by mass spectrometry (ChIP-MS), using either anti-HA antibody-conjugated beads or, as control, IgG-conjugated beads. *Pf*AP2-I was detected exclusively in the HA fraction (Fig. 4a and Supplementary Table 1), indicating that chromatin-bound *Pf*BDP1 interacts with *Pf*AP2-I. In contrast, *Pf*BDP2, *Pf*BDP7 and Chromodomain Protein 1 (*Pf*CHD1), previously reported as *Pf*BDP1 partners^4,10^, were detected by both IP and ChIP-MS (Fig. 4a and Supplementary Table 1), demonstrating that their interaction with *Pf*BDP1 does not depend on chromatin binding. The *Pf*AP2-I and *Pf*BDP1 interaction was further confirmed in the parasite by reciprocal co-immunoprecipitation of *Pf*BDP1-HA with *Pf*AP2-I-GFP, only when we used anti-HA antibody-conjugated beads (Fig. 4b, Supplementary Fig. 4a).

**Fig 4.**
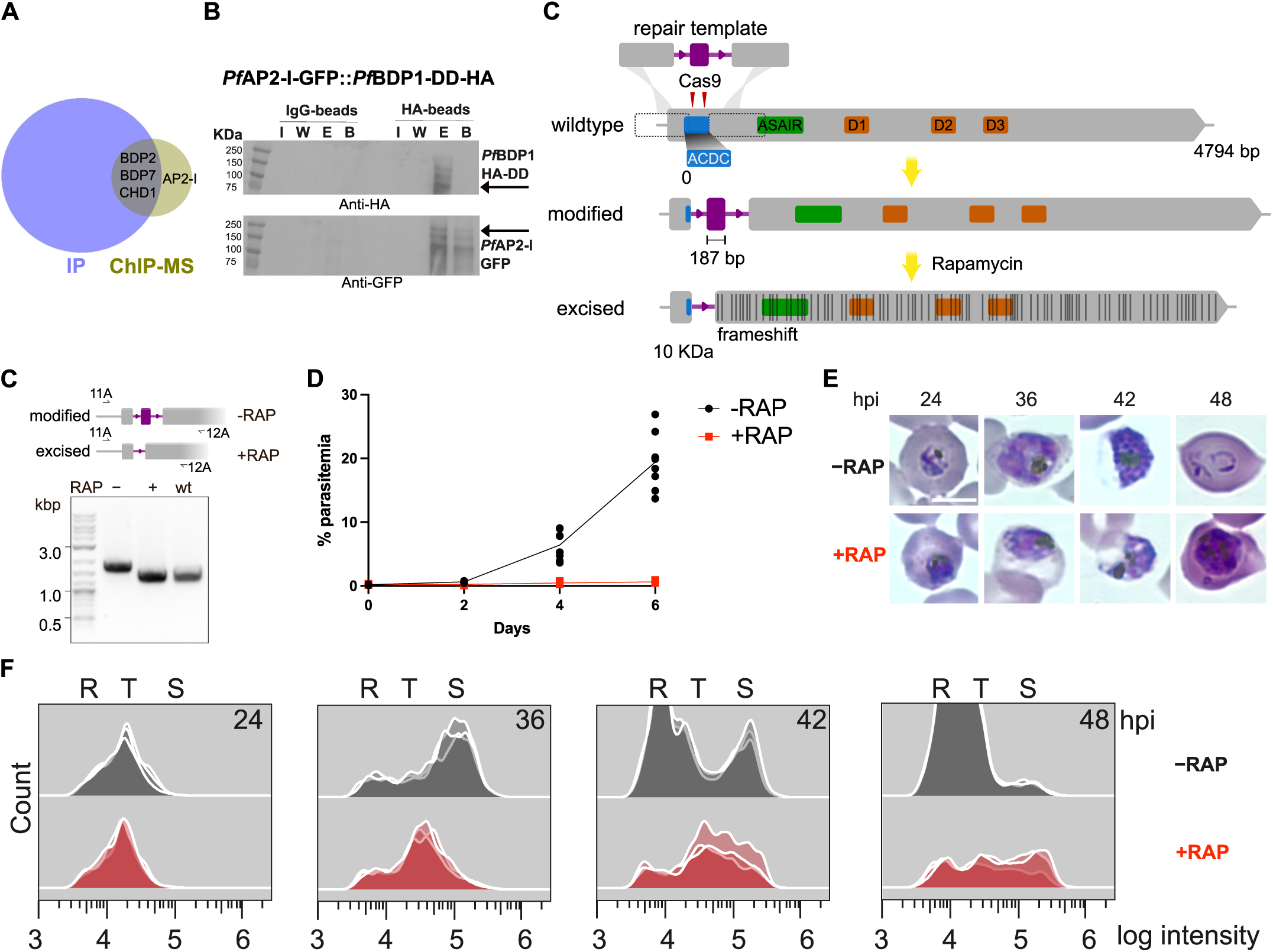
– *Pf*BDP1 associates with *Pf*AP2-I in parasites and *Pf*AP2-I is essential for parasite growth. **a.** The results of immunopurification (IP) versus chromatin immunoprecipitation (ChIP-MS) indicate that only chromatin-bound *Pf*BDP1 interacts with *Pf*AP2-I because this last protein was only identified by ChIP-MS. The number of peptides detected by mass spectrometry is in Supplementary Table 1. **b**. *Pf*AP2-I-GFP is co-immunoprecipitated when *Pf*BDP1-HA-DD is immunoprecipitated with anti-HA conjugated beads, in *Pf*BDP1 iKD parasites expressing endogenously GFP-tagged *Pf*AP2-I (Supplementary Fig. 4a). **c.** A 187 bp region within the ACDC domain (blue) is replaced in the *pfap2-i* locus with a recodonized sequence flanked by a pair of synthetic intron modules (*boxits*), each containing a *loxP* site (arrow head) (Supplementary Fig. 4b). Gene modification is achieved by a Cas9-driven homology directed repair in which double-stranded breaks are introduced in two gRNA target sites (red) simultaneously. Treatment with rapamycin (RAP) induces DiCre-mediated excision of the segment, resulting in multiple downstream frameshift mutations that render the entire downstream section of the gene non-functional. **c**. Diagnostic PCR amplification of the entire modified locus from *ap2i-shiftiko* parasites at 24 h following RAP or mock treatment confirms efficient excision (Locus sequencing in -/+ RAP is in Supplementary Fig. 4c). **d**. Treatment with RAP results in ablation of growth of the *ap2i-shiftiko* parasites. Data shown are from two different growth assays with four biological replicates each. **e.** Representative light microscopic images of Giemsa-stained *ap2i-shiftiko* parasites at specified hours post-invasion (hpi) following RAP (RAP+) and mock (RAP-) treatment at ring stages. RAP-treated mutants are clearly stalled in their development compared to mock-treated controls. Scale bar, 5 μm. **f.** Parasite DNA content was assessed using flow cytometry in SYBR Green-stained RAP- and mock-treated populations. The fluorescence intensity profiles of the AP2I-null parasites (from three independent RAP treatments, Supplementary Fig. 4d) were consistently reduced relative to the control population, indicating developmental arrest at multiple stages of the IDC.

If *Pf*AP2-I recruits *Pf*BDP1 to their target genes, *Pf*AP2-I itself would be expected to be essential. Whilst previous unsuccessful attempts to disrupt *ap2-i* are consistent with this^1,8,9^, AP2-I essentiality was never directly demonstrated. To inducibly disrupt the ∼4.8 kb-long *pfap2-i* gene (Fig. 4c), we used the recently developed frameshift-based trackable inducible knockout (SHIFTiKO) approach^19^. In the *pfap2i-shiftiko* parasites, a 187 bp region, within the ACDC domain, is floxed with a pair of *loxP* synthetic intron modules (Fig. 4c, Supplementary Fig. 4b). DiCre-mediated excision of this region introduces multiple downstream frameshift mutations, rendering the rest of the gene non-functional (Fig. 4c). Treating synchronous ring-stage *pfap2i-shiftiko* parasites with rapamycin (RAP) resulted in efficient excision of the floxed region, 24 hours post-treatment, as confirmed by PCR and sequencing (Fig. 4d, Supplementary Fig. 4c). The RAP-treated *pfap2i-shiftiko* parasites failed to proliferate (Fig. 4d), confirming *Pf*AP2-I’s essentiality for asexual blood stage viability. Detailed phenotypic analyses by microscopy and flow cytometry (Figs. 4e-f, Supplementary Fig. 4d) revealed that *Pf*AP2-I-null mutants growth stalls at the trophozoite stage. Moreover, RAP-treated, schizont-purified *pfap2i-shiftiko* parasites fail to transform into rings (Supplementary Fig. 4e), even when merozoites are allowed to mature inside schizonts, by blocking egress with a protease inhibitor (data not shown), indicating a defect in invasion. Overall, these results indicate that, *Pf*AP2-I plays multiple roles during asexual blood stage development.

### SGC-CBP30 likely binds the *Pf*BDP1 bromodomain

Because interaction of *Pf*BDP1 bromodomain with acetylated histones is not sufficient to recruit *Pf*BDP1 to the chromatin^4^, and BDP1 binding to AP2-I is mediated by the ankyrin and BAAS regions, we investigated whether the *Pf*BDP1 bromodomain is essential for parasite survival. We tested if a truncated *Pf*BDP1 copy containing only the AP2-I interacting domains complements the *Pf*BDP1 iKD^4^. In the iKD parasites, *Pf*BDP1 is fused to a destabilization domain (DD) and it is degraded unless the ligand Shld-1 is added (Supplementary Fig. 5a). Without Shld-1, parasites cannot invade RBCs and die^4^ (Fig. 5a). Expressing only the *Pf*BDP1 BAAS region or the two AP2-I interacting domains (*Pf*BDP1 ANK-BAAS) did not complement the iKD, since parasites were unable to grow in the absence of Shld-1 (Fig. 5a; Supplementary Figs. 5b-c). In contrast, parasites expressing full-length *Pf*BDP1 were viable, regardless of Shld-1 presence (Fig. 5a). These results demonstrate that the *Pf*BDP1 bromodomain is essential for parasite viability.

**Fig 5.**
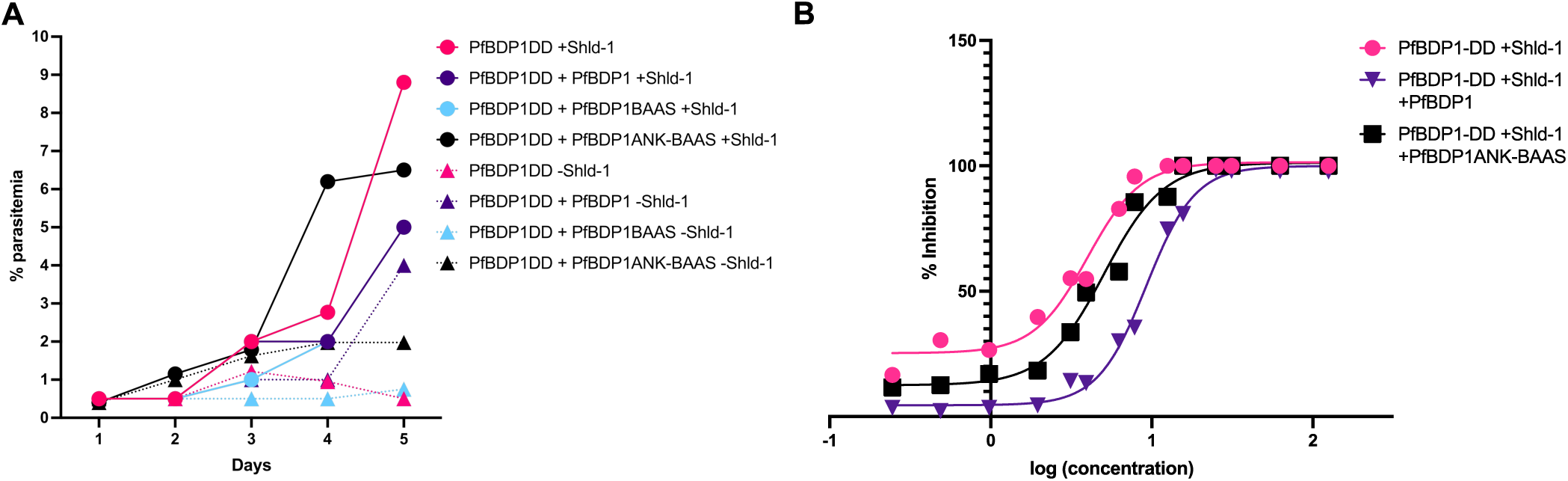
– The *Pf*BDP1 bromodomain is essential and the bromodomain inhibitor SGC-CBP30 *Pf*BDP1 likely targets it. **a.** Parasite survival experiment showing that while *Pf*BDP1 knockdown parasites (pink) die in the absence of Shld-1 (triangle), parasites expressing a second full-length copy of *Pf*BDP1 from a plasmid (*Pf*BDP1DD + *Pf*BDP1, purple) survive with (square) or without Shld-1 (triangle), indicating complementation. On the other hand, parasites expressing from a plasmid just BAAS (*Pf*BDP1DD + *Pf*BDP1BAAS, blue) or the Ankyrin and BAAS (*Pf*BDP1DD + *Pf*BDP1ANK-BAAS, black) die in the absence of Shld-1 (triangle) so the *Pf*BDP1 bromodomain is essential for parasite survival. **b**. Survival assays (n=4) showing a right curve shift, indicative of drug resistance, for parasites overexpressing *Pf*BDP1 (*Pf*BDP1DD+Shld-1 + *Pf*BDP1, purple), as compared to parasites only expressing endogenous *Pf*BDP1 (*Pf*BDP1DD+Shld-1, pink). The same shift is not observed in parasites expressing endogenous *Pf*BDP1 and a second *Pf*BDP1 copy without the bromodomain (*Pf*BDP1DD+Shld-1 + *Pf*BDP1ANK-BAAS, black), suggesting that the probe targets the bromodomain.

We next tested whether the *Pf*BDP1 bromodomain is targeted by the bromodomain inhibitor probe SGC-CBP30, originally designed as an inhibitor of the human CREBB/EP30 bromodomain, as hypothesized^20^. We reasoned that if the compound binds the *Pf*BDP1 bromodomain, overexpression of *Pf*BDP1 should confer resistance by increasing the cellular target pool. To this end, we used the *Pf*BDP1 iKD line expressing a second, episomal, full-length copy of *Pf*BDP1 and cultured the parasites with Shld-1 to maintain expression of both copies. In drug survival assays, the dose-response curve of *Pf*BDP1-overexpressing parasites shifted to the right, relative to that of control parasites expressing only the endogenous *Pf*BDP1 (iKD parasites grown with Shld-1), indicating resistance to SGC-CBP30 (Fig. 5b, Supplementary Fig. 5d). The dose-response curve of parasites expressing a *Pf*BDP1 variant lacking the bromodomain was similar to that of the control (Fig. 5b, Supplementary Fig. 5d), indicating that SGC-CBP30 targets the *Pf*BDP1 bromodomain.

### *Pf*AP2-I and *Pf*BDP1 bind upstream of sexual commitment genes

Recently, 99 genes were identified as markers of sexually-committed *P. falciparum* cells^21^. *Pf*AP2-I and *Pf*BDP1 are expressed in sexually-committed cells^21^ (Supplementary Fig. 6b) and expression of several of these genes is mis-regulated upon *Pf*BDP1 iKD and/or overexpression^4^ (Supplementary Fig. 6a and Supplementary Table 2). Furthermore, among the 99 genes, *Pf*BDP1 binds upstream of 18^4,10^, and *Pf*AP2-I binds upstream of 8^1,15^ (Supplementary Fig. 6a and Supplementary Table 2), including *gdv1* and *ap2-g*, two well-known gametocytogenesis regulators^22,23^. *Pf*AP2-G, activating the sexual differentiation program^22^, has a comparable number of target genes^15^ (Supplementary Fig. 6a and Supplementary Table 2).

*Pf*AP2-G is known to auto-regulate its own expression and to interact with *Pf*AP2-I^15^. To study *Pf*AP2-I binding to the *ap2-g* promoter, independently from *Pf*AP2-G DNA-binding, we used the *P*. *falciparum* F12 strain, expressing a truncated *Pf*AP2-G protein without the AP2 DNA-binding domain (Supplementary Figs. 6c). In F12 parasites, *ap2-g* auto-activation is impaired, yet basal expression persists^24,25^. ChIP of endogenously GFP-tagged *Pf*AP2-I (Supplementary Figs. 6d-e) revealed that *Pf*AP2-I still binds the *ap2-g* promoter in F12 parasites (Fig. 6a). *Pf*AP2-I also binds upstream of *gdv1* (Fig. 6b), not regulated by *Pf*AP2-G, and to the invasion gene *msp4*, but not to *rh4*, consistent with previous findings (Supplementary Fig. 6f). The same results were obtained for *Pf*BDP1 (Figs. 6c-d, Supplementary Fig. 6g). These results suggest that *Pf*BDP1 and *Pf*AP2-I play a role in sexual commitment.

**Fig 6.**
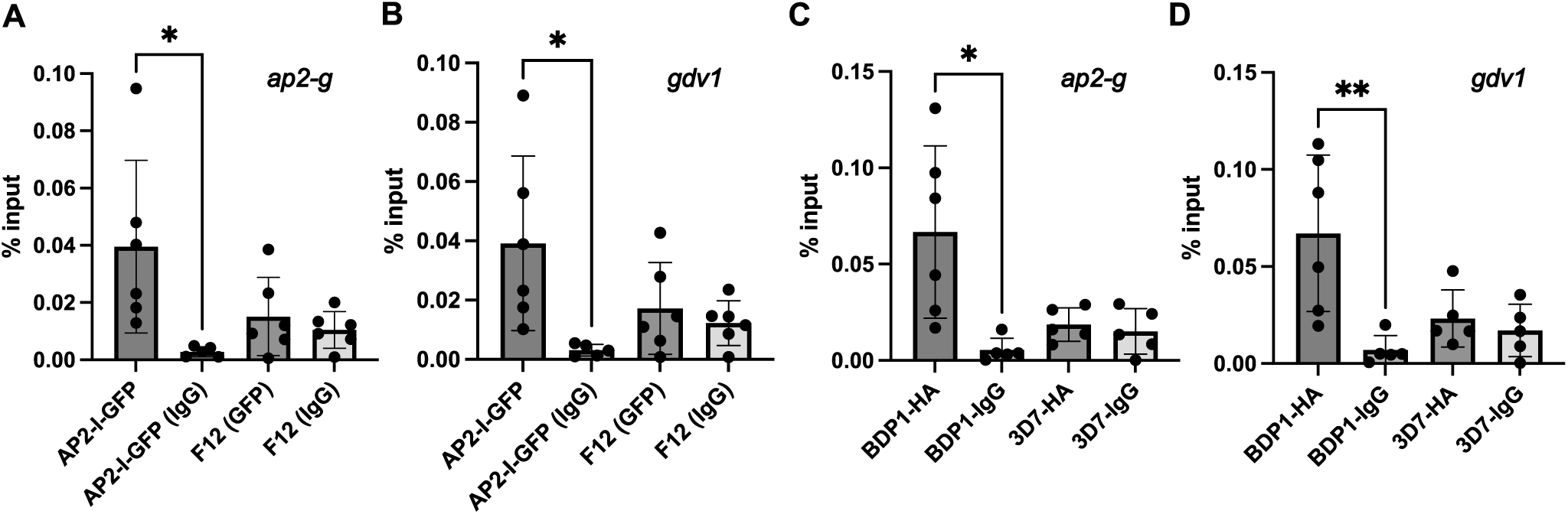
– *Pf*AP2-I and *Pf*BDP1 bind upstream of *ap2-g* and *gdv1*. **a-b.** ChIP-qPCR showing binding of *Pf*AP2-I-GFP expressed in F12 parasites upstream of *ap2-g* (**a**) and *gdv1* (**b**). Because F12 parasites do not express functional *Pf*AP2-G, AP2-I binds independently of *Pf*AP2-G. As control, chromatin was either incubated with IgG rather than anti-GFP, or F12 parasites, not expressing *Pf*AP2-I-GFP, were used. **c-d.** ChIP-qPCR showing binding of *Pf*BDP1-HA upstream of *ap2-g* (**c**) and *gdv1* (**d**). As control, chromatin was either incubated with IgG rather than anti-HA, or 3D7 parasites, not expressing *Pf*BDP1-HA, were used. Results are expressed as mean ± SD, and overall differences between values were evaluated by a two-tailed unpaired t-test. ns, not significant, * P < 0.05.

## Discussion

In this study, we unambiguously demonstrate that AP2-I and BDP1 interact directly. Our *in vitro* experiments reveal that the AP2-I ACDC domain is a key mediator of this interaction, thereby assigning for the first time a functional role to this specific domain of the ApiAP2 proteins. This result is consistent with our recent structural analysis, which suggested that the ACDC acts as a protein-protein interaction domain^26^. It is likely that in other ApiAP2 proteins, the ACDC mediates interaction with other chromatin-related proteins.

The AP2-I ACDC binds to a conserved region of BDP1 that we call BAAS. Within this sequence, we identify a predicted α-helix that may be directly involved in this interaction. We also show that the BDP1 ankyrin domain, a well-known protein-protein interaction scaffold, binds a highly conserved region of AP2-I that we call ASAIR. ASAIR is restricted to *Plasmodium* AP2-I orthologues and it is only the second conserved region described in ApiAP2 proteins, beyond the AP2 domains, after the ACDC.

Although we could not determine the precise stoichiometry of the AP2-I/BDP1 complex, our data indicate that complex formation promotes AP2-I dimerization. Alternatively, AP2-I may dimerize upon chromatin binding, since its DNA-binding motif in RBC stages frequently occurs in tandem within gene promoters^1,18,27^.

Supporting a functional role for the AP2-I/BDP1 complex in a chromatin context, the interaction between the BDP1 ankyrin domain and the AP2-I ASAIR region is compatible with DNA-binding. Our results in *P*. *falciparum* further suggest that *Pf*BDP1 is recruited to the chromatin through interaction with *Pf*AP2-I bound to its specific DNA-binding motif in target promoters. Since *Pf*BDP1 is also present at loci not targeted by *Pf*AP2-I^1,4,10^, it may be recruited at these sites *via* another, as-yet unidentified, DNA-binding protein. However, no additional ApiAP2 proteins were detected by ChIP-MS.

We show that the *Pf*BDP1 bromodomain is essential for parasite viability, underscoring the importance of its interaction with acetylated histones. This likely fine-tunes *Pf*BDP1 recruitment to specific loci and/or supports its role in chromatin remodeling, consistent with other bromodomain-containing proteins^28^. We further demonstrate that overexpression of *Pf*BDP1 increases resistance to the bromodomain inhibitor SGC-CBP30, and that this effect is dependent on the presence of the bromodomain, suggesting that SGC-CBP30 binds the *Pf*BDP1 bromodomain. However, while bromodomain inhibitors kill parasites^20,29^, the large number of human bromodomain proteins raises the risk of off-target effects. Targeting instead the Apicomplexa-specific BAAS sequence, would more selectively inhibit BDP1.

Unlike the *Pf*BDP1 knockdown parasites that are defective in invasion but grow normally in RBCs^4^, the growth of the *Pf*AP2-I-null mutants is impaired, indicating that *Pf*AP2-I plays multiple roles during the intraerythrocytic developmental cycle (IDC), perhaps by activating transcription of different sets of genes. Further experiments are required to test this and to determine if these other functions require interaction with *Pf*BDP1.

We show that both *Pf*AP2-I and *Pf*BDP1 bind upstream of *gdv1* and *ap2-g*, and *Pf*AP2-I binding is independent of interaction with *Pf*AP2-G, consistent with earlier observations^15^. Unfortunately, we could not evaluate the number of sexually-committed cells in the *Pf*AP2-I null mutants or the *Pf*BDP1 iKD because the parent strains are poor gametocyte producers^22,30^.

The functional interplay between *Pf*AP2-I and *Pf*AP2-G, and whether *Pf*BDP1 associates with the *Pf*AP2-I/*Pf*AP2-G complex or it dissociates from the *ap2-g* promoter, upon *Pf*AP2-G binding, remains unclear. Although *Pf*AP2-G was not detected by ChIP-MS, its expression is limited to a small cell fraction, and most genes co-bound by *Pf*AP2-I and *Pf*AP2-G^15^ are also bound by *Pf*BDP1^4,10^ (Supplementary Table 3), keeping this question open.

Overall, our results establish the essential role of the AP2-I/BDP1 complex in *P. falciparum* and highlight both proteins as attractive drug targets. We identify two novel interacting regions - BAAS, an Apicomplexa-specific region of BDP1, and ASAIR, a *Plasmodium*-specific region of AP2-I – and assign a protein-protein interaction function to the Apicomplexa-specific ACDC domain of AP2-I. Each of these regions could be exploited for selective inhibition. While targeting BAAS or ASAIR would specifically inhibit BDP1 or AP2-I, respectively, targeting ACDC could inhibit multiple ACDC-containing ApiAP2 proteins. Since these are interaction domains, such inhibitors would also likely block complex formation. Because the two proteins are expressed across developmental stages and regulate multiple processes, their inhibitors may also have broad efficacy throughout the lifecycle.

## Methods

### Optimized genes synthesis

Codon-optimized full-length *ap2-I* and *bdp1* from *P. vivax* were synthesized (Thermo Fisher Scientific) to reduce the risk of poor protein expression. A C-terminal 6xHis-Tag was added to the *Pv*BDP1 construct, while a C-terminal Strep-Tag®II was added to the *Pv*AP2-I construct to facilitate protein purification.

The recodonised sequence of *pfap2-i* and the flanking *boxit* modules (*boxit-*:RR:*boxit+*) used to generate the null-mutant parasites were synthesized commercially by Azenta Life Sciences.

### Cloning and expression in insect cells

Synthetic genes were amplified using primers 1 and 470 or 2 and 1396 (Supplementary Table 4), and cloned into the Multibac baculovirus/insect cell expression vector system. The full-length *Pv*AP2-I gene was inserted by In-Fusion cloning (Takara bio) between the Tn7R and Tn7L sites of the pFL donor plasmid, under the control of the constitutive polyhedrin promoter, using BamHI and EcoRI restriction sites. For co-expression, the *Pv*BDP1 gene was cloned into the mini-attTn7 transposition acceptor site, under the control of the constitutive P10 promoter, using XhoI and NcoI restriction enzymes.

The resulting pFL plasmids were introduced into a yellow fluorescent protein-containing bacmid by heat shock transformation into EmBacY *Escherichia coli* DH10 cells. Recombinant EmBacY bacmids were purified using the Monarch® Plasmid Miniprep Kit (New England Biolabs) according to the manufacturer’s instructions, and subsequently transfected into Sf21 insect cells with X-tremeGENE HP DNA transfection reagent (Roche) to generate first-generation (V0) virus.

Following amplification, viral stocks were titrated by serial dilution, using yellow fluorescent protein expression as a marker of infection. For protein production, Sf21 cells were infected with baculovirus at a multiplicity of infection (MOI) of 5 × 10^−3^ Pfu/cell and harvested 5–6 days post-infection (3–4 days after proliferation arrest). Cells were sonicated, and the supernatant containing the second-generation virus (V1) was collected, titrated, supplemented with 5% serum to prevent aggregation, and stored at 4°C in the dark.

For large scale protein expression, 600 mL cultures of Sf21 cells at 0.5 x 10^6^ cells/mL were infected with V1 virus at a final MOI of 5 x 10^-3^ Pfu/cell. Infected cells were harvested, the supernatant discarded, and cell pellets containing intracellular recombinant proteins were frozen in liquid nitrogen and stored at -20°C.

### Cloning and expression in bacteria

The synthetic full-length *Pv*AP2-I gene, C-terminally tagged with a Strep-tag II, was amplified using primers 429/432 (Supplementary Table 4) and inserted into the NcoI/XhoI sites of a pET28a(+) vector (Novagen), which carries a kanamycin resistance cassette. The synthetic full-length *Pv*BDP1 gene, C-terminally tagged with a 6xHis tag, was amplified using primers 402/403 (Supplementary Table 4) and cloned into the NdeI/XhoI sites of a pET21a(+) vector (Novagen), carrying an ampicillin resistance cassette.

Gene fragments of *Pv*AP2-I and *Pv*BDP1 were PCR-amplified using the primers listed in Supplementary Table 4 and cloned into either pET28a(+) or pET21a(+) vectors (Novagen), respectively. Cloning was performed using T4 DNA ligase (Thermo Fisher Scientific) and the restriction enzyme pairs (Thermo Fisher Scientific) listed in Supplementary Table 4. All *Pv*AP2-I constructs were designed with a C-terminal Strep-Tag II, while *Pv*BDP1 fragments carried a C-terminal 6xHis tag.

Plasmids were first transformed into ultra-competent *E. coli* DH5α cells (Thermo Fisher Scientific) and correct assembly was confirmed by PCR using Taq 2X Master Mix (New England BioLabs) followed by Sanger sequencing (Genewiz).

For protein expression, verified plasmids were transformed into *E. coli* (DE3)-Gold cells. Cultures were grown in 2xYT medium at 37°C for 3 h, induced with 0.5 mM IPTG (Sigma), and harvested by centrifugation. Cell pellets were flash-frozen in liquid nitrogen and stored at -20°C until use.

### Protein purification

Bacterial or insect cell pellets were resuspended in buffer A (20 mM Tris-HCl, pH 7-8 adjusted according of the isoelectric point (pHi) of the protein, 200 mM NaCl) supplemented with a protease inhibitor cocktail (Thermo Fisher Scientific). Cells were lysed by sonication using a probe-tip sonicator (Branson). Soluble fractions were subsequently purified by affinity chromatography.

Strep-tagged *Pv*AP2-I and its fragments were purified using a StrepTrap HP column (Cytiva) equilibrated in buffer A at 4°C on an FPLC system. After extensive washing with buffer A, bound Strep-tagged proteins were eluted with buffer A supplemented with 2.5 mM desthiobiotin. This procedure was also applied to samples containing His-tagged *Pv*BDP1 or its fragments (either co-expressed or pre-purified and mixed) to assess interaction with immobilized Strep-tagged *Pv*AP2-I.

His-tagged *Pv*BDP1 and its fragments were purified using Ni-NTA affinity chromatography on benchtop columns. After thorough washing with cold buffer A, bound proteins were eluted with increasing imidazole concentrations (up to 400 mM) in cold buffer A.

For large-scale purification, size-exclusion chromatography (SEC) was performed as a second purification step using either Hiload 16/60 Superdex S75 or Superdex S200 prep grade columns (GE Healthcare), equilibrated in buffer A, depending on the molecular weight of the target protein or fragment. Each SEC column was calibrated using a Gel Filtration Calibration Kit (Cytiva).

Protein concentrations were determined by measuring absorbance at 280nm using a NanoDrop spectrophotometer (Thermo Fisher Scientific). Purified proteins were concentrated using Vivaspin concentrators, aliquoted, and stored at -80°C in buffer A.

### Electrophoretic Mobility Super-shift Assays

Experiments were performed using the *Pv*AP2-I (157-510) fragment, which includes the ASAIR region and the first AP2 DNA-binding domain of the protein. The DNA substrate was a double-stranded 5’-Cy5-labeled oligonucleotide (Eurofins Genomics), derived from the promoter region of the PVP01_0824200 gene, encoding an uncharacterized *P. vivax* protein (the primers sequence is in Supplementary Table 4). This 60 bp sequence contains the GTCGAC motif, previously identified as the AP2-I D1 domain binding site^18^.

To assess DNA-binding activity, increasing concentrations of the *Pv*AP2-I fragment (10 to 1280 nM) were incubated with 3 nM of the labeled DNA for 30 min on ice in binding buffer (20 mM Tris-HCl pH 7.5, 50 mM NaCl, 2 mM DTT, 0.05% Nonidet P40). This allowed determination of the minimum protein concentration required to observe a DNA shift. To test potential cooperative binding, increasing concentrations (160 to 1280 nM) of the *Pv*BDP1 ankyrin domain (residues 1–220) were added to a fixed concentration of *Pv*AP2-I (320 nM) and were incubated with the DNA probe under the same conditions. Samples were resolved on an 8% native polyacrylamide gel, and the super-shifts of the fluorescent DNA probe were visualized using a Typhoon™ RGB Imaging System (Cytiva). Control experiments confirmed that the ankyrin domain alone did not bind to the DNA probe under these conditions.

### Mass photometry analysis

To assess the molecular mass and infer the quaternary structure of the purified proteins and protein complex, mass photometry experiments were performed at room temperature using an automated TwoMP mass photometer (Refeyn). Protein samples were analyzed in purification buffer A at concentrations ranging from 10 to 100 nM. Prior to sample measurements, the instrument was calibrated using Bovine Serum Albumin (BSA, 66kDa) and Immunoglobulin G (IgG, 150kDa) standards. The molecular mass of individual particles landing on the measurement surface was determined based on the interference between the light scattered by the molecule and the light reflected by the measurement surface. The signal measured is called the interferometric contrast and is directly correlated with molecular mass^31^.

### Dynamic Light Scattering

Dynamic Light Scattering (DLS) was used to assess the size of the proteins and detect potential conformational changes upon complex formation. Experiments were carried out at 25°C using a Prometheus Panta intrument (NanoTemper Technologies) with high-sensitivity capillaries. Protein samples were analyzed at a concentration of 20 µM with a laser excitation set to 54% at 405 nm. The diffusion coefficient of each sample was determined, and the corresponding hydrodynamic radius (Rh) was calculated by using the Stokes-Einstein equation, as implemented in Panta Analysis software (NanoTemper).

### Thermal shift assays

Fluorescence-based thermofluor experiments were performed at 25°C in 96-well plates on a StepOne qPCR instrument (Thermo Fisher Scientific) using a final protein concentration ranging from 20 to 40 µM, depending on the proteins and fragments, in buffer B (20 mM Tris-HCl, pH 7-8 adjusted to match the pHi of the target protein, 100 mM NaCl). The temperature was gradually increased from 25°C to 95°C at a rate of 0.5 °C/min in the presence of 1x Sypro® Orange dye. Fluorescence was monitored at 570 nm as the dye bound to hydrophobic regions exposed during protein unfolding. The “PTS clear plate” setting was used as the background calibration reference. Melting curves were analyzed using the manufacturer’s software, and the melting temperature (Tm) of the protein was determined by plotting the derivative of fluorescence intensity as a function of temperature.

### Cloning of parasite plasmids

Genomic DNA from *P. falciparum* parasites was extracted using the DNeasy Kit (Qiagen) following the manufacturer’s instructions.

To express *Pf*BDP1 in the parasite, the full-length gene was amplified from parasite genomic DNA using primers 311 and 352 (Supplementary Table 4). The ankyrin and BAAS domains of *Pf*BDP1 were amplified from genomic DNA using primers 532/534 and 238/533, respectively, and assembled by Gibson reaction. All resulting PCR fragments were inserted into the pUF1-mRuby-NLS plasmid, which had been previously digested with MluI and XhoI. The pUF1-mRuby-NLS plasmid was generated by amplifying mRuby from a synthetic mRuby-NLS fragment (IDT) using primers 182/183, followed by cloning into the pUF1 vector^32^, digested with MluI and PacI.

To generate the CRISPR/Cas9 plasmids for GFP-tagging the endogenous *ap2-I* gene in F12 parasites and *Pf*BDP1 knockdown parasites, the guide RNA (gRNA) cassette was constructed by annealing primers 273/274 (Supplementary Table 4) and cloning into the BbsI sites of the pDC2-Cas9-gRNA plasmid^33^ via Golden Gate assembly. The donor plasmid was generated by cloning a first homology region corresponding to the 3’ end of *ap2-I* (amplified with primers 36/37, Supplementary Table 4) into the pBCAM-GFP plasmid^1^, previously digested with BglII and NotI. A second homology region corresponding to the gene 3’untranslated region (UTR) was amplified using primers 158/159 and inserted by Gibson assembly.

To inducibly knockout *ap2-I*, the SHIFTiKO strategy (Ramaprasad & Blackman 2024) was employed as follows. Two high-quality gRNA sequences in close proximity (<200 bp) to each other and upstream or within the ACDC domain were chosen from gRNAs predicted using EuPaGDT. A region that spans these two Cas9 cleavage sites and of length not divisible by three (187 bp) was chosen as the target segment to be recodonized and floxed (RR). This recodonised sequence and the flanking *boxit* modules (*boxit-*:RR:*boxit+*) was synthesized commercially (Azenta Life Sciences). To assemble the donor plasmid (pSKO-ap2i), a 532 bp long synthesized segment was amplified using a common primer pair, boxit.F and boxit.R (Supplementary Table 4), and the two ∼480 bp long homology arms was amplified from parasite genomic DNA with primers carrying common 20 bp extensions (Supplementary Table 4). Column-purified amplicons were then assembled into a standard backbone vector in a modular one-pot In-Fusion (Takara) reaction, according to kit instructions. The targeting plasmid (pCas9Duo-ap2i) was assembled as follows. gRNA sequences were ordered as oligos with unique pairs of overhangs (ATTG and AAAC for gRNA1 and TTGG and TAAA for gRNA2, Supplementary Table 4), annealed and inserted into the dual-guide targeting plasmid, pCas9-Duo^19^ in a simple one-pot Golden Gate assembly. Successful insertion of gRNA1 and gRNA2 was confirmed by Sanger sequencing using M13.R and M13.F_R primers respectively.

All plasmids were fully sequence-verified by Nanopore sequencing (Plasmidsaurus and Eurofins).

### Parasite culture and transfection

*P. falciparum* parasites were cultured at 37°C under an atmosphere of 5% oxygen and 5% carbon dioxide in RPMI 1640 media (Gibco) supplemented with sodium hypoxanthine (0.002%), 0.5% Albumax II (Invitrogen) and gentamycin (10 ug/mL), and maintained at 2% hematocrit, unless stated otherwise. *Pf*BDP1 knockdown parasites were additionally supplemented with 0.5µM Shld-1 (Ozyme), unless stated otherwise, and 2.5nM WR99210 (WR) (Jacobs Pharmaceuticals). The DiCre-expressing *P. falciparum* B11 line^30^ and *Pf*AP2-I conditional knockout parasites were maintained at 37°C in human red blood cells (NHS Scotland) in HEPES-buffered RPMI 1640 supplemented with 0.5% Albumax II, Glutamine (0.03%), Glucose (0.2%), sodium hypoxanthine (0.002%) and gentamycin (10 ug/mL) and gassed with 5% CO2 and 1% O2. Parasite cultures were fed with fresh blood every other day, and parasitemia was maintained between 2% and 5%, unless stated otherwise, as assessed by counting the number of parasites in blood smears.

Human blood was obtained from the French Blood Institute (Etablissement Français du Sang, EFS). Plasmid transfections into *P. falciparum* parasites were carried out by electroporation using a Gene-Pulser II system (Bio-Rad), as previously described^34^. Briefly, ring-stage parasites at 5% hematocrit were resuspended to 50% in cytomix buffer (120 mM KCl, 0.2 mM CaCl_2_, 2 mM EGTA, 10 mM MgCl_2_, 25 mM Hepes, 5 mM K_2_HPO_4_, 5 mM KH_2_PO_4_, pH 7.6) and electroporated with 100μg of plasmid DNA, or 30µg of each CRISPR plasmid. Following electroporation, parasites were resuspended in warm complete media containing 2% hematocrit and incubated overnight. The next day, transfectants were selected using 1.5 μM DSM1 (5-methyl[1,2,4]triazolo[1,5-a]pyrimidin-7-yl)naphthalen-2-ylamine; Sigma) and 2.5 μg/ml blasticidin-S-HCl (BSD). F12 and BDP1 knockdowns CRISPR parasites stopped being drug-selected once GFP integration into the desired locus was PCR-confirmed.

Alternatively, to generate the *ap2i-shiftiko* inducible knockout line was generated by isolating mature schizonts for either transfections or routine synchronization of parasite cultures were done as described previously ^35^ by centrifugation over 73% (v/v) isotonic Percoll (GE Healthcare, Life Sciences) cushions. Transfection was performed by introducing 20 μg of pCas9Duo-ap2i and 60 μg of linearised pSKO-*ap2-i* (purified using PureYield™ Plasmid Midiprep Kit, Promega) into 20 µL Percoll-enriched schizonts by electroporation using an Amaxa 4D Nucleofector X (Lonza), using program FP158 as previously described ^36^. Drug selection with 2.5 nM WR99210 was applied 24 h post-transfection for 4 days with successfully transfected parasites arising 2 weeks post-transfection. As previously shown ^19^, with SHIFTiKO, transfection yielded near-homogenous population of modified *ap2i-shiftiko* parasites so transfected parasites were used directly for subsequent phenotyping without establishing a clonal line. To obtain AP2I-null parasites, DiCre-mediated excision of the target floxed region in synchronous early ring-stage *ap2i*-*shiftiko* parasites (2 hr post-invasion) was induced by rapamycin treatment (100 nM RAP for 3 hr or 10 nM overnight). DMSO-treated parasites were used as wildtype controls.

Parasite synchronization to the ring stage was performed as previously described^37^ by incubating pelleted cultures 1:1 with warm 0.3M L-Alanine, 10mM Hepes pH 7.5 solution, for 10 min at 37°C, followed by centrifugation and resuspension of the pellet in fresh culture medium. Routine synchronization of parasite culture of the *ap2i-shiftiko* inducible knockout lines was done as described previously ^35^ by centrifugation over 73% (v/v) isotonic Percoll (GE Healthcare, Life Sciences) cushions. Then, isolated schizonts were let to rupture and invade fresh erythrocytes for 2 h at 120 rpm, followed by removal of residual schizonts by another Percoll separation and sorbitol treatment to finally obtain a highly synchronized preparation of newly invaded ring-stage parasites.

To induce *Pf*BDP1 knockdown, synchronized cultures were washed to remove Shld-1 from the medium.

### IC50 drug assays

Drug sensitivity assays were performed as previously described in^38^. Briefly, synchronized *P. falciparum* cultures were adjusted to 1.75% parasitemia and 2% hematocrit, and assays were conducted in duplicate in 96-well plates. Cultures were incubated at 37°C for 72 hours in the presence of serial dilutions of the drug. After incubation, parasites were lysed by freeze-thaw cycles and transferred to a new 96-well black detection plate. SYBR® Green I nucleic acid stain (Thermo Fisher Scientific) was added, and plates were incubated in the dark for 3 hours at room temperature. Fluorescence was then measured at excitation and emission wavelengths of 587 and 610 nm, respectively, using a Tecan spectrophotometer (Tecan). Dose-response curves and half-maximal inhibitory concentration (IC_50_) values were calculated from log-transformed fluorescence counts using GraphPad Prism (GraphPad Software).

### ChIP-qCPR

Chromatin immunoprecipitation (ChIP) was performed as previously described in^1^. Briefly, 40 hours schizont-stage parasite cultures were harvested, lysed with saponin, and formaldehyde-crosslinked. The isolated chromatin was extracted and sheared in SDS lysis buffer to generate a DNA fragment of 100-150 bp, using an S220 focused-ultrasonicator (Covaris) with the following settings: peak power 105W, 2% duty factor, 200 cycles per burst, and total treatment time of 80 seconds. The chromatin extracts were then pre-cleared with ChIP-grade Protein A+G magnetic beads (Thermo Fisher Scientific). An input control sample was retained, and the remaining chromatin was incubated overnight at 4°C with 1 μg of anti-GFP (Abcam Ab290) or anti-HA (Roche) or, as a negative control, the same amount of IgG (Diagenode). Immuno-complexes were captured with Protein A+G magnetic beads, washed extensively, and eluted using an elution buffer (1% SDS, 0.1M NaHCO_3_). The input and ChIP samples were reverse cross-linked overnight at 45°C in the presence of 0.4M NaCl and purified by phenol:chloroform extraction. The concentration of each eluted sample was determined using a Nanodrop spectrophotometer (Thermo Fisher Scientific), followed by 1:10 dilution, and the samples were used for quantitative PCR (qPCR) reactions performed in triplicate wells using Sso FastGreen (ThermoFisher Scientific). Each ChIP-qPCR experiment was performed with at least 3 biological replicates. ChIP-qPCR data were analyzed using the ΔΔCt method. Primer sequences used for qPCR are listed in Supplementary Table 4.

### IP-MS and ChIP-MS

The IP was performed as previously described in^1^. For ChIP-MS, chromatin was collected as for ChIP, but then the sheared chromatin was diluted 1:10 in dilution buffer (20mM HEPES pH7.9, 1mM EDTA, 1mM EGTA, 40% glycerol, protease inhibitors). IP and ChIP-MS samples were incubated for 90 minutes with Dynabeads M270-Epoxy beads (Thermo Fisher Scientific) conjugated with anti-HA antibodies (Roche) or control IgG, as described in ^1^.

Eluted samples were mixed with the BCA reactif in Laemli buffer and protein reduction and alkylation was carried on, followed by protein digestion using the single-pot treatment, solid-phase-enhanced sample-preparation (SP3) approach as described in ^39^. Digestion was performed overnight at 37°C using 5 ng/µl of sequencing-grade modified trypsin (Promega). Trypsin-generated peptides were dried, resuspended in loading buffer (2% acetonitrile, 0.05% Trifluoroacetic acid) and 5 µl were analyzed by nanoLC-MSMS using a nanoElute liquid chromatography system (Bruker) coupled to a timsTOF Pro2 mass spectrometer (Bruker). MS and MS/MS spectra were recorded over a range of *m/z* 100 to 1700 with an ion mobility scan range from 0.75-1.25 V.s/cm^2^. Data were acquired in parallel accumulation – serial fragmentation (PASEF) ion mobility-based acquisition mode, with 10 PASEF MSMS scans per cycle.

Raw MS and MS/MS data were processed and converted into .mgf files using Bruker DataAnalysis software. Protein identification was performed using the Mascot search engine (Matrix Science, London, UK) against a *Plasmodium falciparum* database downloaded from Uniprot (20210715), including common contaminants such as trypsin and streptavidin. Trypsin was set as the digestion enzyme, allowing up to two miscleavages. Carbamidomethylation of cysteines weas set as a fixed modification and oxidation of methionines as a variable modification. Peptide and fragment mass tolerances were set to 15 ppm and 0.05 Da, respectively.

### Western blot analysis and Fluorescent gels

For Western blot analyses, proteins were transferred from SDS-PAGE gels onto 0.45 µm nitrocellulose membranes (Thermo Fischer Scientific). Detection of *Pv*AP2-I and its fragments was performed using a murine monoclonal anti-Strep-Tag®II primary antibody (StrepMAB-Classic, IBA), followed by an HRP-conjugated goat anti-mouse IgG secondary antibody (Thermo Fisher Scientific). *Pv*BDP1 and its fragments were detected directly using an HRP-conjugated anti-6xHis antibody (Merck). The presence of the antigen was revealed using the colorimetric substrate TMB (3,3‘,5,5’-Tetramethylbenzidine, Thermo Fisher Scientific), which undergoes a color change upon HRP-mediated oxidation. Developed blots were photographed for archiving.

For *P. falciparum* parasite samples, nuclear fractionation was performed as described in^39^. mRuby- or GFP-tagged proteins were detected using a mouse anti-mRuby primary antibody (St John) or a mouse anti-GFP primary antibody (Roche) at a 1:1000 dilution, respectively. Histone 3 was detected using a 1:1000 dilution of a mouse anti-H3 antibody (Abcam). Detection was achieved using an HRP-conjugated goat anti-mouse secondary antibody (Fisher) at a 1:3000 dilution. HA-tagged proteins were detected using a 1:1500 dilution of a mouse anti-HA HRP-conjugated antibody (Cell Signaling). Bound antibodies were visualized using the ECL Plus chemiluminescent substrate (Pierce), and images were acquired using a ChemiDoc imaging system (BioRad). Alternatively, mRuby proteins were detected directly in SDS-Page gels with a Typhoon imager (Cytiva) using the 532 nm laser with the Cy3 filter.

### Microscopy and Flow cytometry assays

To stain with DAPI, magnetically purified schizonts^40^, using LD columns (Miltenyi Biotec), were fixed in 1XPBS/4%formaldehyde, washed and deposited into glass slides mounted with VectaShield Antifade medium containing DAPI (Vector laboratories) and imaged with a SP8 confocal microscope (Leica) at 63x amplification. Images were treated with ImageJ.

### Growth and development profiling of the *ap2-i* conditional knockout parasites

Growth assays were performed to assess parasite growth across 3–4 erythrocytic replication cycles. Synchronous cultures of ring-stage *ap2i-shiftiko* parasites at 0.1% parasitaemia and 2% haematocrit were maintained as 3-4 replicates per treatment condition (RAP or DMSO) in 12-well plates. 50 µL from each well was sampled at 0, 2, 4, and 6 days post-RAP treatment, fixed with 50 µL of 0.2% glutaraldehyde in 1XPBS and stored at 4°C for flow cytometry quantification. Fixed parasites were stained with SYBR Green (Thermo Fisher Scientific, 1:10,000 dilution) for 20 min at 37°C and analyzed by flow cytometry on a BD FACS Celesta. For every sample, parasitaemia was estimated by recording 10,000 events and filtering with appropriate forward and side scatter parameters and gating for SYBR Green stain-positive (infected RBCs) and negative RBCs using a 527/32 detector configuration. All data were analyzed using FlowJo software.

Parasite development was monitored by microscopic examination at selected timepoints using Giemsa-stained thin blood films. Samples were also fixed at these timepoints for flow cytometry analysis. Fluorescence intensity of the SYBR Green stain-positive population was quantified to assess DNA content, the increase of which was taken as a proxy for growth stage progression.

To test for invasion, 44h percoll-purified schizonts, pre-treated or not for 4 hours with the C2 egress inhibitor, which blocks egress reversibly^41^, were incubated with fresh red blood cells, shaking, at 37°C, for 4 hours, in the presence or absence of RAP. The parasitemia was counted at the beginning of the assay and after 4 hours. The percentage of schizonts or rings at the end of the 4 hours was determined by Giemsa-stained smears.

### Bioinformatics

The protein schemes of AP2-I and BDP1 were drawn to scale using the tool MyDomain from Prosite (https://prosite.expasy.org/mydomains/).

Multiple sequence alignments (MSA) were performed using ClustalW^42^ on the PRABI^43^ platform. Secondary structure elements were added to the top of MSA using ESPript^44^. Secondary structures were predicted using PSIPRED^45^. Three-dimensional structural models of proteins and individual domains were predicted using AlphaFold3^46^ with default settings. Model quality was assessed using predicted Template Modeling scores (pTM) and interface pTM scores (ipTM). Reliable predictions were defined as those with pTM values > 0.5 and ipTM values > 0.8, in accordance with established criteria^47,48^. Predicted Aligned Error (PAE) plots, which provide an estimate of the positional and orientational error between residue pairs, were plotted for further analysis of the proposed models. In these 2D plots, low-error regions (i.e., confident predictions) appear as dark green, light green areas indicate less reliable inter-domain positioning. For protein complexes, additional boxes representing inter-chain predictions were included.

Final structural models were visualized and analyzed using the PyMOL Molecular Graphics System, Version 3.1.1^49^. The AlphaFold3 models were colored according to the pLDDT (predicted Local Distance Difference Test), a confidence score ranging from 0 (orange) to 100 (blue) indicating the most reliable regions of the proposed model.

### Phylogenetic trees

Protein sequences were aligned using Clustal Omega^50^ and the resulting alignment was processed with Jalview^51^ to build the phylogenetic trees.

### Antibodies

Mouse anti-GFP (Roche): AB_390913

Mouse anti-histone H3 (Abcam): AB_470239

Rat anti-HA high-affinity 3F10 (Roche): AB_10094468

Rabbit anti-GFP (Abcam): AB_302295

Rabbit anti-IgG (Abcam): AB_2614925

Mouse anti-IgG (Abcam): AB_2827163

Strep-mAb classic (IBA): AB_513133

HRP-conjugated anti-6xHis antibody (Merck): AB_10947552

Mouse anti-mRuby primary antibody (St John): STJ140251

Mouse anti-HA HRP-conjugated antibody (Cell Signaling): AB_1264166

## Acknowledgements

This work benefited from the facilities and expertise of the I2BC platforms (Gif-sur-Yvette, France) supported by IBiSA, FRISBI, Ile de France Region, Plan Cancer, CNRS and Paris-Saclay University: the proteomic platform (Proteomic-Gif, SICaPS) and more particularly Laïla Sago, the Macromolecular Interaction Measurements platform and the Imagerie-Gif core facility supported by the Agence Nationale de la Recherche (ANR-10-INBS-04/FranceBioImaging ; ANR-11-IDEX-0003-02/ Saclay Plant Sciences). We would also like to acknowledge Flow Core facility within the MVLS Shared Research Facilities, University of Glasgow.

This work received funding from the French Proteomic Infrastructure (ProFI; attributed to JMS), MICROBES Paris-Saclay University Interdisciplinary Object (attributed to SN), HEALTHI Paris-Saclay University Interdisciplinary Object (attributed to JMS and SN) and the ATIP-Avenir CNRS program (attributed to JMS). PT was funded by a DIM1Health Ile de France region grant (attributed to JMS). AR and ARS are funded by the UKRI MRC Career Development Award (MR/Z504762/1).

We thank https://PlasmoDB.org and https://OrthoMCL.org for invaluable resources. We are grateful to Bjorn Kafsack for kindly providing the F12 parasite strain; Sebastian Baumgarten for the pDC2-Cas9-gRNA plasmid; Daan Noordemeer for the AlexaFluor secondary antibody; and Michael Duffy and Michaela Petter for the BDP1 knockdown parasite strain.

3D7 parasites were obtained through BEI Resources, NIAID, NIH: *Plasmodium falciparum*, Strain 3D7, MRA-102, contributed by Daniel J. Carucci. The polyclonal anti-*Plasmodium falciparum* Pfs16 antiserum (Rabbit), MRA-1276, was also obtained through BEI Resources, NIAID, NIH, contributed by Kim C. Williamson.

## Author contributions

JMS and SN conceptualized the project. JMS, SN and AR designed the experiments. MLB performed the *in vitro* experiments. LM, together with PT, conducted parasite experiments. SA carried out the EMSA assays. VR assisted with protein expression in insect cells. MC performed the phylogenetic analysis. CR generated the *Pf*AP2-I-GFP(F12) strain. ARS generated and analyzed the *Pf*AP2-I conditional KO parasites. JMS, SN, MLB, LM, AR, ARS and PT analyzed the data. JMS and SN wrote the manuscript, acquired funding and supervised the project.

## Supplementary information

**Supplementary Table 1**

List of proteins identified by IP-MS and ChIP-MS.

**Supplementary Table 2**

List of genes upregulated in sexually committed cells, as determined by Dogga *et al*., that are ChIP-Seq target genes of AP2-I, AP2-G and BPD1, as determined by Santos *et al*. and Josling *et al*..

**Supplementary Table 3**

List of genes that are ChIP-Seq target genes of *Pf*AP2-G, *Pf*AP2-I and *Pf*BDP1 as determined by Josling *et al*.

**Supplementary Table 4**

List of primers used in this study.

---

**Supplementary Figure 1.**
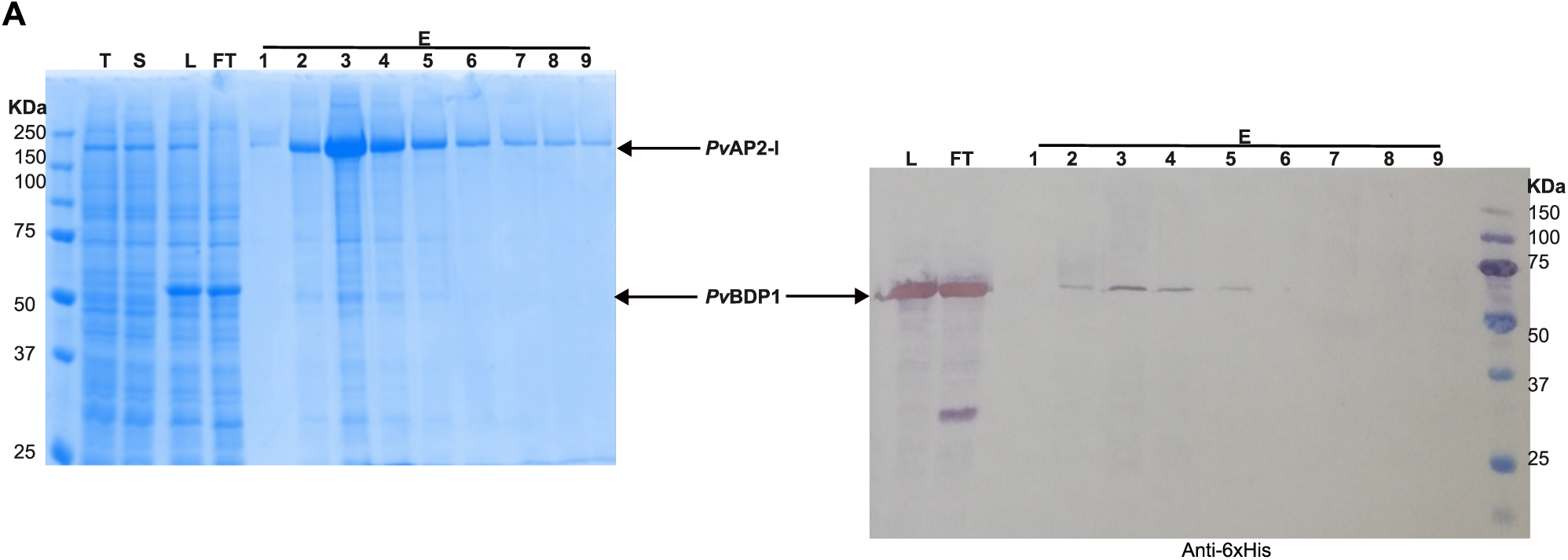
Co-purification of Strep-tagged *Pv*AP2-I and His-tagged *Pv*BDP1. His-tagged *Pv*BDP1 (55kDa) expressed *in E. coli* was added to the insect cell lysate containing Strep-tagged *Pv*AP2-I (151 kDa) prior to loading on a Strep-Trap column. **a.** SDS-PAGE analysis. **b.** Western blot analysis using anti-6xHis antibodies (because *Pv*AP2-I is clearly detected by SDS-PAGE, we did not confirm its presence by western-blot). Samples: T (total insect cell extract), S1 (soluble fraction of insect cells), L: loaded mixture of S and *Pv*BDP1, FT: flow-through, 1-9: elution fractions.

**Supplementary Figure 2.**
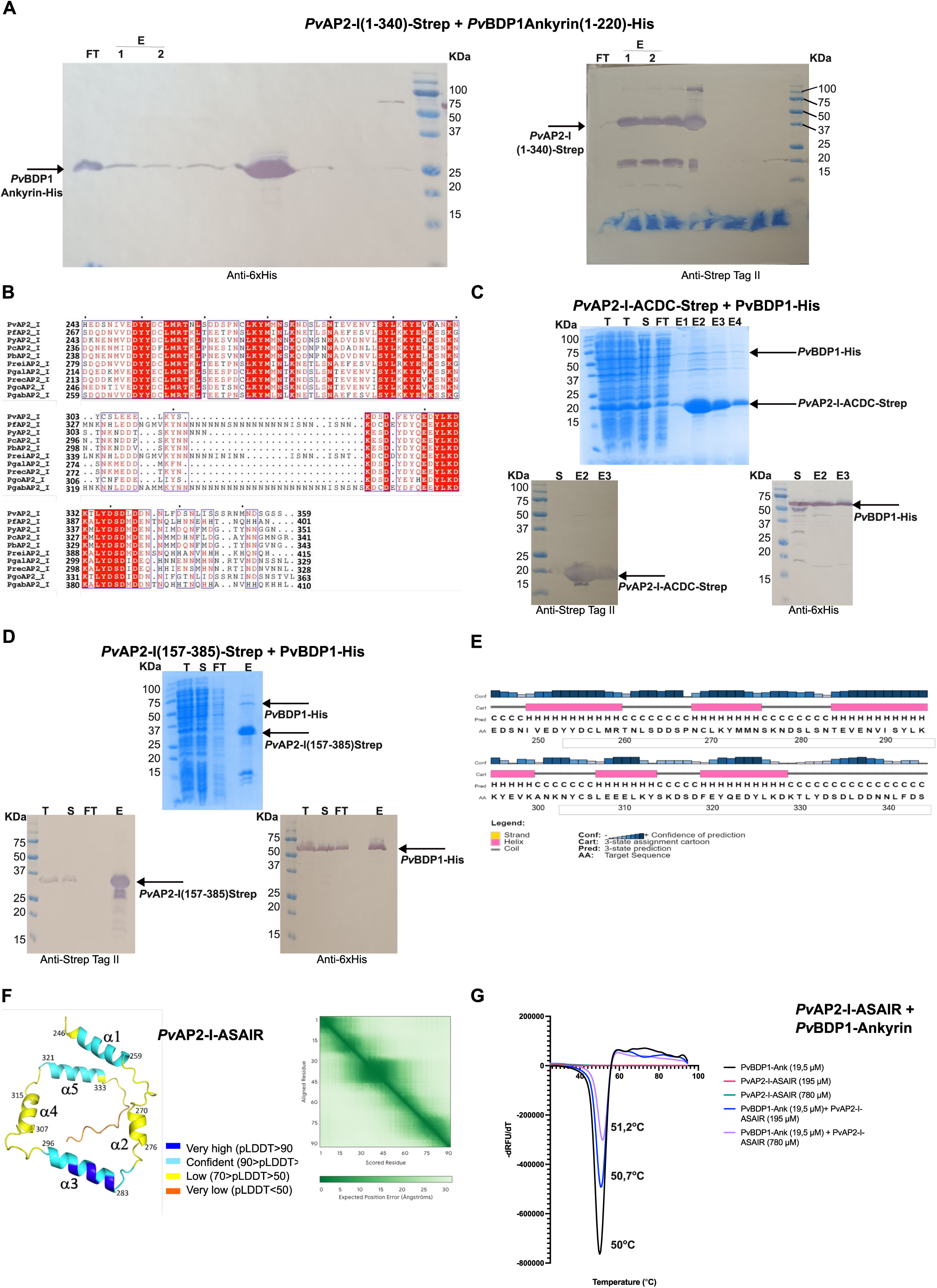
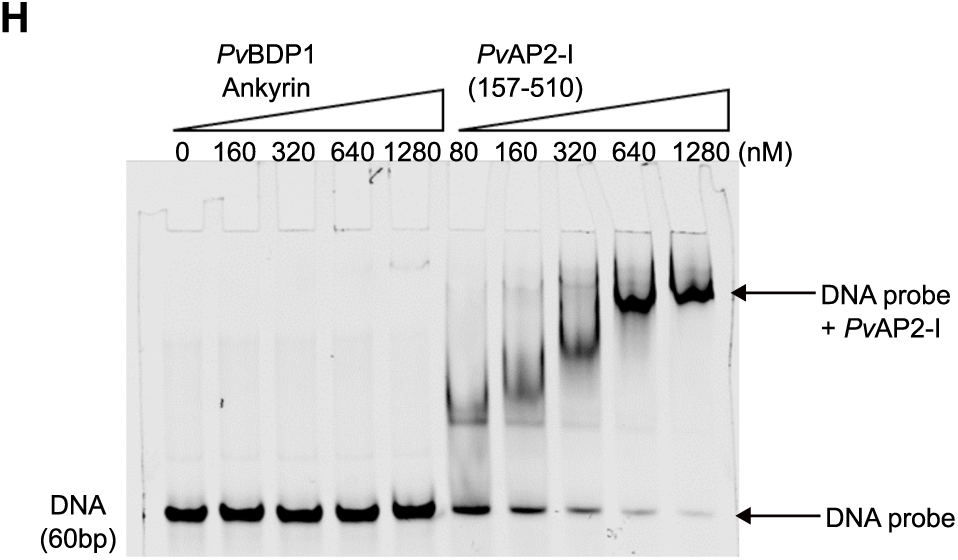
Bioinformatic and biochemical analyses of the AP2-I conserved sequence. **a.** Non-cropped images of the western-blots shown on figure 2b. **b**. Multiple sequence alignment of the conserved region of *Pv*AP2-I (*P. vivax* A0A564ZT73), *Pf*AP2-I (*P. falciparum* Q8IJW6), *Py*AP2-I (*P. yoelii* A0AAE9WZ09), *Pc*AP2-I (*P. chabaudi* A0A4V0K907), *Pb*AP2-I (*P. berghei* A0A509ATM1), *Prei*AP2-I (*P. reichnowi* A0A2P9DFA7), *Pgal*AP2-I (*P. gallinaceum* A0A1J1GPA6), *Prec*AP2-I (*P. rectilum* A0A1J1H3Z6), *Pgo*AP2-I (*P. gonderi* A0A1Y1JE14) and *Pgab*AP2-I (*P. gaboni* A0A151LKI2). Residues are colored according to their conservation level: white characters on a red background indicate strictly conserved residues across all aligned sequences; red characters indicate partial conservation; black characters indicate no significant conservation; blue frames highlight cluster of conserved residues. Asterisks above the alignment mark every 10 residues in the reference sequence (*Pv*AP2-I). **c-d**. Co-purification on a Strep-trap column of full-length His-tagged *Pv*BDP1 co-expressed in *E. coli* with Strep-tagged fragments of *Pv*AP2-I containing either the isolated ACDC domain (residues 30-180, 18 kDa) (**c**) or the adjacent conserved sequence (residues 157-385, 26 kDa) (**d**). SDS-PAGE and Western blot show co-elution between *Pv*BDP1 and the two fragments. Samples: T (total cell extract), S (soluble fraction), FT (flow-through), E (concentrated elution fraction). **e**. Secondary structure prediction of the *Pv*AP2-I ASAIR region (residues 250-340) using PsiPred. Prediction confidence is shown in blue, predicted β-strands in yellow, α-helices in pink and coils in grey. **f**. AlphaFold3 rank 1 model of *Pv*AP2-I ASAIR. The 3D structure is colored by predicted Local Distance Difference Test (pLDDT) confidence scores. The accompanying Predicted Aligned Error (PAE, right) plot illustrates the poor reliability of the overall 3D fold. **g**. Effect of the *Pv*AP2-I ASAIR fragment containing helix α3 (residues 280-310) on the thermal stability of the ankyrin domain of *Pv*BDP1 (residues 1-220). Thermal unfolding was monitored by plotting the first derivative of the relative fluorescence units as a function of temperature (−dRFU/dT). The melting temperature (*T_m_*) corresponds to the minimum of each curve. Black: BDP1 ankyrin domain alone (19.5 µM), green: AP2-I ASAIR alone (780µM), blue: mixture of *Pv*BDP1 ankyrin domain (19.5µM) with AP2-I ASAIR at 195µM, purple: mixture of *Pv*BDP1 ankyrin domain (19.5µM) with AP2-I ASAIR at 780µM. **h**. Electro Mobility Shift Assay (EMSA) showing that the *Pv*BDP1 ankyrin domain alone does not bind DNA, unlike the *Pv*AP2-I fragment (157-510) encompassing ASAIR and the first AP2 domain.

**Supplementary Figure 3.**
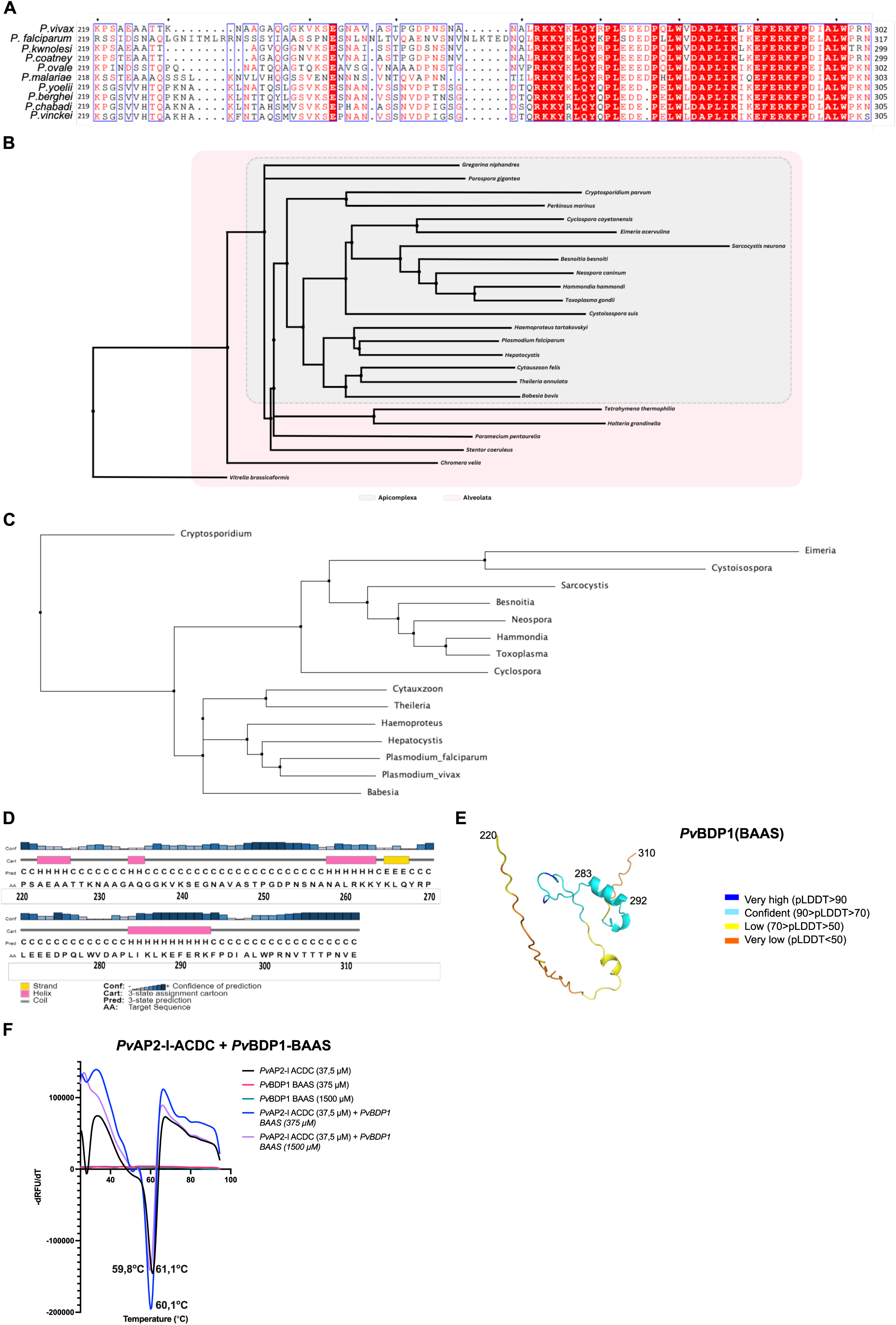
Bioinformatic and functional analyses of the central region of *Pv*BDP1. **a**. Multiple sequence alignment of the central region (residues 220-310) of *Pv*BDP1 (*P. vivax* A0A8S4HEK5), *Pf*AP2-I (*P. falciparum* Q8IJW6), *Pk*BDP1 (*P. knowlesi* XP_002261856), *Pco*BDP1 (*P. coatney* XP_019913821), *Po*BDP1 (*P. ovale* SBT76169), *Pm*BDP1 (*P. malariae* XP_028862223), *Py*BDP1 (*P. yoelii* XP_022811693), *Pb*BDP1 (*P. berghei* XP_034420416), *Pc*BDP1 (*P. chabaudi* SCL99649) and *Pv*iBDP1 (*P. vinckei* CAD2099431). Residues are color-coded according to their conservation level: white characters on a red background indicate strictly conserved residues across all aligned sequences; red characters indicate partial conservation; black characters indicate no significant conservation; blue frames highlight cluster of conserved residues. Asterisks above the alignment mark indicate every 10 residues in the reference sequence (*Pv*AP2-I). **b**. Phylogenetic tree of BDP1 proteins showing that some Alveolata encode BDP1-like proteins characterized by the presence of both an ankyrin domain and a bromodomain. Alveolata clades are highlighted in pink. **c**. Phylogenetic tree of the BDP1 proteins containing a BAAS-like sequence showing that only the BPD1 proteins from Apicomplexa contain a BAAS region. The sequence alignment used to build the tree is shown. **d**. Secondary structure prediction of the *Pv*BDP1 BAAS region using PsiPred. Prediction confidence is shown in blue, predicted β-strands in yellow, α-helices in pink and coils in grey. **e**. AlphaFold3 model of the *Pv*BDP1 BAAS region (residues 220-310). The 3D structure is colored by predicted Local Distance Difference Test (pLDDT) confidence scores. **f**. Effect of the *Pv*BDP1 BAAS peptide (residues 275-296) on the thermal stability of the *Pv*AP2-I fragment (30-180) containing the ACDC domain. Melting temperatures (*T_m_*) were determined by plotting the first derivative of relative fluorescence units as a function of temperature (−dRFU/dT). *T_m_* values correspond to the minima of the curves. Black: AP2-I ACDC domain alone (37,5µM), pink: BDP1 BAAS alone (375µM), green: BDP1 BAAS alone (1500µM), blue: mixture of AP2-I ACDC domain (37.5µM) with BDP1 BAAS peptide at 375µM), and purple: mixture of AP2-I ACDC domain (37.5µM) with BDP1 BAAS peptide at 1500µM).

**Supplementary Fig 4.**
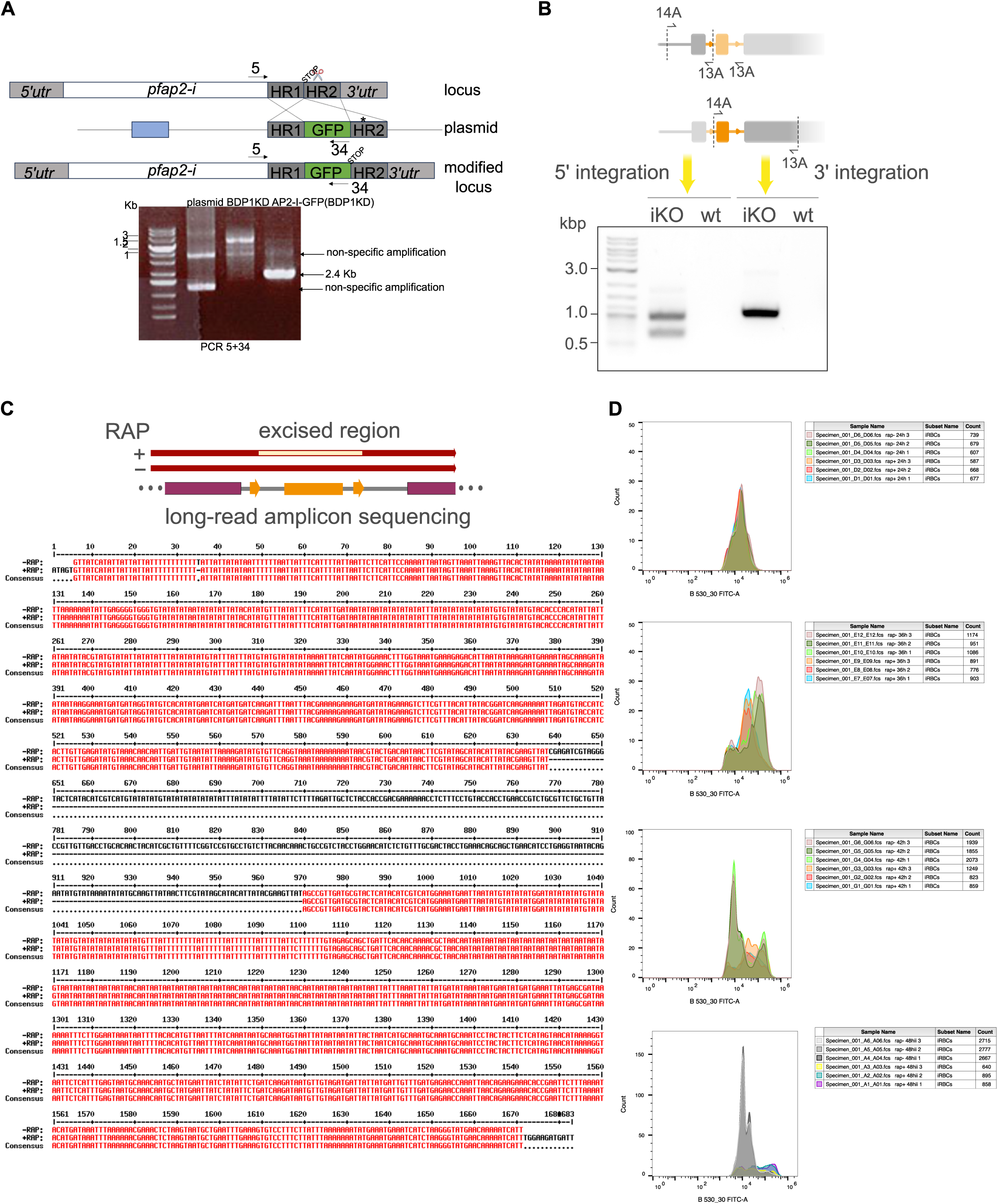
*Pf*BDP1 cKD parasites expressing *Pf*AP2-I-GFP and *Pf*AP2-I conditional knockout parasites. **a.** Diagnostic PCR shows successful integration of the GFP cassette in the *pfap2-i* locus only in the transfected parasites AP2-I-GFP(BDP1-KD) and not in the parental strain (BDP1 KD). The primers sequence is in Supplementary Table 4. **b**. Diagnostic PCR shows successful integration of the repair sequence in the *pfap2-i* locus. Outer primers (11A and 12A) and standard inner primers (13A and 14A) were used to check for 5’ and 3’ integration. 13A has two binding sites within the repair sequence, thus yielding two PCR products for 5’ integration. The primers sequence is in Supplementary Table 4. **c**. Long-read sequencing of the amplicons of the modified and excised locus show precise excision of the floxed region in the RAP-treated parasites. **d**. Non-processed flow cytometry data at different times post-invasion showing growth stall in the *pfap2-i*-null mutants only in the presence of RAP. **e**. Invasion assays indicate that, after 4-hour incubation with red blood cells, non-treated parasites (gray) invade host cells and transform into rings (ring.mia and parasitemia are equal), whereas RAP-treated parasites (red) remain as schizonts (sch.mia and parasitemia are very similar and the ring.mia value is very low).

**Supplementary Fig 5.**
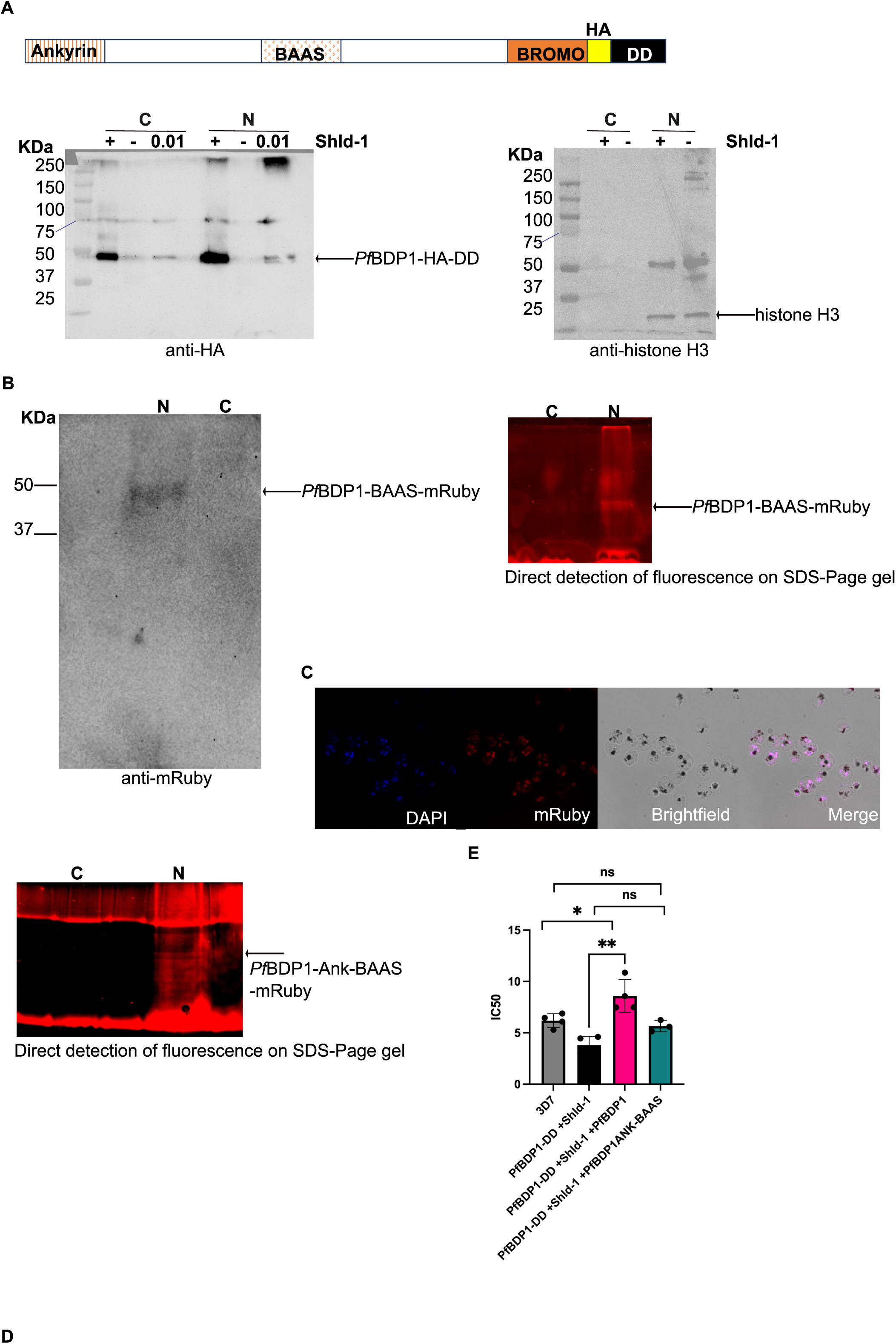
Study of *Pf*BDP1 bromodomain. **a**. Western-blot showing that *Pf*BDP1 is knockdown in the absence of Shld-1. Histone H3 was used as a loading control. - : no Shld-1, +: 0.5µM Shld-1, 0.01: 0.01 µM Shld-1. **b**. Western-blot and SDS-Page gel showing expression of *Pf*BDP1-BAAS-mRuby in the nucleus of the *Pf*BDP1 knockdown parasites. **c**. Imaging of *Pf*BDP1-BAAS-mRuby parasites showing overlap between the DAPI and the mRuby staining, indicating that the protein is nuclear. **d**. SDS-Page gel showing expression of *Pf*BDP1-Ank-BAAS-mRuby in the nucleus of the *Pf*BDP1 knockdown parasites as determined by the detection of a fluorescent signal corresponding to the mRuby fusion protein only in the nuclear fraction. The detection of multiple bands might indicate that there is protein degradation. In **a**, **b** and **c**: C: cytosolic fraction, N: nuclear fraction. **d.** Bar graph showing that the IC50 to SGC-CBP30 of parasites over-expressing PfBDP1 (*Pf*BDP1-DD + Shld-1 + *Pf*BD1, pink) is higher than that of 3D7 parasites (parental line, gray), *Pf*BDP1 knockdown parasites expressing endogenous *Pf*BDP1 (*Pf*BDP1-DD + Shld-1, black) or of *Pf*BDP1 knockdown parasites expressing endogenous *Pf*BDP1 and a protein variant without the bromodomain (*Pf*BDP1-DD + Shld-1 + *Pf*BDP1-Ank-BAAS, green). Results are expressed as mean ± SD, and overall differences between values were evaluated by a two-tailed unpaired t-test. ns, not significant, * P < 0.05, ** P < 0.005.

**Supplementary Fig 6.**
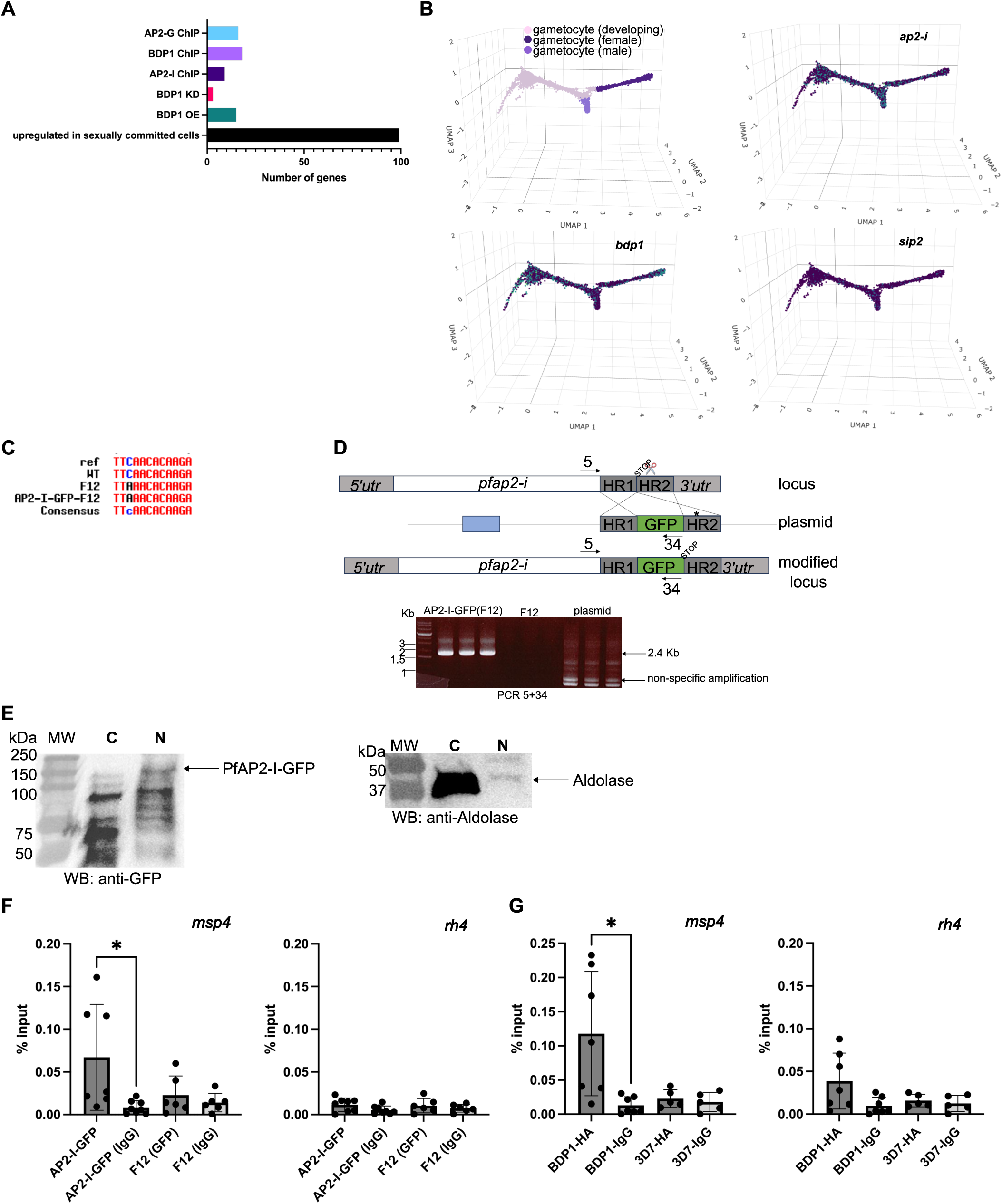
Role of *Pf*AP2-I and *Pf*BDP1 in sexual commitment. **a.** Number of genes overexpressed in sexually committed cells (black), as determined by Dogga *et al*., that are mis-regulated upon *Pf*BDP1 overexpression (OE, green) or knockdown (KD, pink) or that are ChIP-Seq targets of *Pf*AP2-I (dark purple), *Pf*BDP1 (light purple) or *Pf*AP2-G (blue), according to Santos *et al*. and Josling *et al*.. **b.** Snapshots from the malaria cell atlas (https://www.malariacellatlas.org/atlas/plasmodium-falciparum-atlas/) showing that *Pf*AP2-I and *Pf*BDP1, but not *Pf*SIP2, are expressed by sexually committed cells (developing gametocytes) according to Dogga et al.. In light blue are cells expressing the transcripts. **c.** Sanger sequencing indicates that the mutation carried by F12 parasites leading to a premature STOP codon in the *ap2-g* locus is present in the transgenic parasites *Pf*AP2-I-GFP(F12). WT are 3D7 parasites. Ref is the sequence from the genomic database PlasmoDB. **d.** Scheme of the *ap2-I* locus before and after introduction of the GFP-tag in C-terminus by CRISPR-Cas9. A scissor indicates the Cas9 cleavage site. The position of the STOP codon as well as the position of the primers used for PCR diagnosis are indicated. Amplification of a 2.4 Kb product from genomic DNA of the transgenic parasites *Pf*AP2-I-GFP(F12) indicates integration of the GFP tag. **e.** Western-blot showing expression of *Pf*AP2-I-GFP in the nucleus (N) of the transgenic parasites *Pf*AP2-I-GFP(F12). On the other hand, aldolase is only expressed in the cytosolic (C) fraction. **f**. ChIP-qPCR showing that AP2-I-GFP binds to the *msp4* promoter but not to the *rh4* one. **g**. ChIP-qPCR showing that BDP1-HA binds to the *msp4* promoter but not to the *rh4* one. As control, chromatin was either incubated with IgG rather than anti-GFP/anti-HA or F12/3D7 parasites, not expressing *Pf*AP2-I-GFP/*Pf*BDP1-HA, were used.

## References

1. Santos, J. M. et al. Red Blood Cell Invasion by the Malaria Parasite Is Coordinated by the PfAP2-I Transcription Factor. Cell Host Microbe 21, 731–741.e10 (2017).

2. Balaji, S., Babu, M. M., Iyer, L. M. & Aravind, L. Discovery of the principal specific transcription factors of Apicomplexa and their implication for the evolution of the AP2-integrase DNA binding domains. Nucleic Acids Res 33, 3994–4006 (2005).

3. Oehring, S. C. et al. Organellar proteomics reveals hundreds of novel nuclear proteins in the malaria parasite Plasmodium falciparum. Genome Biol 13, R108 (2012).

4. Josling, G. A. et al. A Plasmodium Falciparum Bromodomain Protein Regulates Invasion Gene Expression. Cell Host Microbe 17, 741–751 (2015).

5. Singh, A. K. et al. Structural insights into acetylated histone ligand recognition by the BDP1 bromodomain of Plasmodium falciparum. Int J Biol Macromol 223, 316–326 (2022).

6. Hoeijmakers, W. A. M. et al. Epigenetic reader complexes of the human malaria parasite, Plasmodium falciparum. Nucleic Acids Res 47, 11574–11588 (2019).

7. Zhang, M. et al. Uncovering the essential genes of the human malaria parasite Plasmodium falciparum by saturation mutagenesis. Science 360, eaap7847 (2018).

8. Modrzynska, K. et al. A Knockout Screen of ApiAP2 Genes Reveals Networks of Interacting Transcriptional Regulators Controlling the Plasmodium Life Cycle. Cell Host Microbe 21, 11–22 (2017).

9. Shang, X. et al. Genome-wide landscape of ApiAP2 transcription factors reveals a heterochromatin-associated regulatory network during Plasmodium falciparum blood-stage development. Nucleic Acids Res 50, 3413–3431 (2022).

10. Quinn, J. E. et al. The Putative Bromodomain Protein PfBDP7 of the Human Malaria Parasite Plasmodium Falciparum Cooperates With PfBDP1 in the Silencing of Variant Surface Antigen Expression. Front Cell Dev Biol 10, 816558 (2022).

11. Toenhake, C. G. et al. Chromatin Accessibility-Based Characterization of the Gene Regulatory Network Underlying Plasmodium falciparum Blood-Stage Development. Cell Host Microbe 23, 557–569.e9 (2018).

12. Lindner, S. E. et al. Total and putative surface proteomics of malaria parasite salivary gland sporozoites. Mol Cell Proteomics 12, 1127–1143 (2013).

13. López-Barragán, M. J. et al. Directional gene expression and antisense transcripts in sexual and asexual stages of Plasmodium falciparum. BMC Genomics 12, 587 (2011).

14. Silvestrini, F. et al. Protein export marks the early phase of gametocytogenesis of the human malaria parasite Plasmodium falciparum. Mol Cell Proteomics 9, 1437–1448 (2010).

15. Josling, G. A. et al. Dissecting the role of PfAP2-G in malaria gametocytogenesis. Nat Commun 11, 1503 (2020).

16. Moreno-Pérez, D. A., Dégano, R., Ibarrola, N., Muro, A. & Patarroyo, M. A. Determining the Plasmodium vivax VCG-1 strain blood stage proteome. J Proteomics 113, 268–280 (2015).

17. Russell, T. J. et al. Inhibitors of ApiAP2 protein DNA binding exhibit multistage activity against Plasmodium parasites. PLoS Pathog 18, e1010887 (2022).

18. Campbell, T. L., Silva, E. K. D., Olszewski, K. L., Elemento, O. & Llinás, M. Identification and Genome-Wide Prediction of DNA Binding Specificities for the ApiAP2 Family of Regulators from the Malaria Parasite. PLOS Pathogens 6, e1001165 (2010).

19. Ramaprasad, A. & Blackman, M. J. A scaleable inducible knockout system for studying essential gene function in the malaria parasite. Nucleic Acids Res 53, gkae1274 (2025).

20. Chua, M. J., Robaa, D., Skinner-Adams, T. S., Sippl, W. & Andrews, K. T. Activity of bromodomain protein inhibitors/binders against asexual-stage *Plasmodium falciparum* parasites. International Journal for Parasitology: Drugs and Drug Resistance 8, 189–193 (2018).

21. Dogga, S. K. et al. A single cell atlas of sexual development in Plasmodium falciparum. Science 384, eadj4088 (2024).

22. Kafsack, B. F. C. et al. A transcriptional switch underlies commitment to sexual development in malaria parasites. Nature 507, 248–252 (2014).

23. Filarsky, M. et al. GDV1 induces sexual commitment of malaria parasites by antagonizing HP1-dependent gene silencing. Science 359, 1259–1263 (2018).

24. Painter, H. J., Carrasquilla, M. & Llinás, M. Capturing in vivo RNA transcriptional dynamics from the malaria parasite Plasmodium falciparum. Genome Res 27, 1074–1086 (2017).

25. Bancells, C. et al. Revisiting the initial steps of sexual development in the malaria parasite Plasmodium falciparum. Nat Microbiol 4, 144–154 (2019).

26. Le Berre, M. et al. Structural characterization of the ACDC domain from ApiAP2 proteins, a potential molecular target against apicomplexan parasites. Acta Cryst D 81, 38–48 (2025).

27. Young, J. A. et al. In silico discovery of transcription regulatory elements in Plasmodium falciparum. BMC Genomics 9, 70 (2008).

28. Josling, G. A., Selvarajah, S. A., Petter, M. & Duffy, M. F. The Role of Bromodomain Proteins in Regulating Gene Expression. Genes (Basel) 3, 320–343 (2012).

29. Amann, M. et al. A Novel Inhibitor against the Bromodomain Protein 1 of the Malaria Pathogen Plasmodium Falciparum. ChemMedChem 20, e202500024 (2025).

30. The Actinomyosin Motor Drives Malaria Parasite Red Blood Cell Invasion but Not Egress | mBio. https://journals-asm-org.insb.bib.cnrs.fr/doi/full/10.1128/mbio.00905-18.

31. Young, G. et al. Quantitative mass imaging of single biological macromolecules. Science 360, 423–427 (2018).

32. Ghorbal, M. et al. Genome editing in the human malaria parasite Plasmodium falciparum using the CRISPR-Cas9 system. Nat Biotechnol 32, 819–821 (2014).

33. Mussgnug, S. et al. N6-methyladenosine primes the malaria parasite for transmission. 2025.03.28.645929 Preprint at 10.1101/2025.03.28.645929 (2025).

34. Fidock, D. A. & Wellems, T. E. Transformation with human dihydrofolate reductase renders malaria parasites insensitive to WR99210 but does not affect the intrinsic activity of proguanil. Proc Natl Acad Sci U S A 94, 10931–10936 (1997).

35. Harris, P. K. et al. Molecular Identification of a Malaria Merozoite Surface Sheddase. PLOS Pathogens 1, e29 (2005).

36. Moon, R. W. et al. Adaptation of the genetically tractable malaria pathogen Plasmodium knowlesi to continuous culture in human erythrocytes. Proceedings of the National Academy of Sciences 110, 531–536 (2013).

37. Braun-Breton, C., Rosenberry, T. L. & da Silva, L. P. Induction of the proteolytic activity of a membrane protein in Plasmodium falciparum by phosphatidyl inositol-specific phospholipase C. Nature 332, 457–459 (1988).

38. Nkhoma, S. C. et al. Dissection of haplotype-specific drug response phenotypes in multiclonal malaria isolates. Int J Parasitol Drugs Drug Resist 15, 152–161 (2021).

39. Hughes, C. S. et al. Single-pot, solid-phase-enhanced sample preparation for proteomics experiments. Nat Protoc 14, 68–85 (2019).

40. Boyle, M. J. et al. Isolation of viable Plasmodium falciparum merozoites to define erythrocyte invasion events and advance vaccine and drug development. Proc Natl Acad Sci U S A 107, 14378–14383 (2010).

41. Collins, C. R. et al. Malaria parasite cGMP-dependent protein kinase regulates blood stage merozoite secretory organelle discharge and egress. PLoS Pathog 9, e1003344 (2013).

42. Clustal W: improving the sensitivity of progressive multiple sequence alignment through sequence weighting, position-specific gap penalties and weight matrix choice | Nucleic Acids Research | Oxford Academic. https://academic.oup.com/nar/article/22/22/4673/2400290.

43. Combet, C., Blanchet, C., Geourjon, C. & Deléage, G. NPS@: Network Protein Sequence Analysis. Trends in Biochemical Sciences 25, 147–150 (2000).

44. Gouet, P., Courcelle, E., Stuart, D. I. & M√©toz, F. ESPript: analysis of multiple sequence alignments in PostScript. Bioinformatics 15, 305–308 (1999).

45. McGuffin, L. J., Bryson, K. & Jones, D. T. The PSIPRED protein structure prediction server. Bioinformatics 16, 404–405 (2000).

46. Jumper, J., et al. Highly accurate protein structure prediction with AlphaFold. Nature 596, 583–589 (2021).

47. Xu, J. & Zhang, Y. How significant is a protein structure similarity with TM-score = 0.5? Bioinformatics 26, 889–895 (2010).

48. Abramson, J., et al. Accurate structure prediction of biomolecular interactions with AlphaFold 3. Nature 630, 493–500 (2024).

49. Schrödinger, L. & DeLano, W. PyMOL. (2020).

50. Madeira, F. et al. The EMBL-EBI Job Dispatcher sequence analysis tools framework in 2024. Nucleic Acids Res 52, W521–W525 (2024).

51. Waterhouse, A. M., Procter, J. B., Martin, D. M. A., Clamp, M. & Barton, G. J. Jalview Version 2—a multiple sequence alignment editor and analysis workbench. Bioinformatics 25, 1189–1191 (2009).

